# A bifunctional H/ACA snoRNP mediates both pseudouridylation and rRNA scaffolding during ribosome assembly

**DOI:** 10.64898/2026.03.20.713187

**Authors:** Jutta Hafner, Matthias Thoms, Hussein Hamze, Anna Forstner, Ismaël Alidou-D’Anjou, Hafiza Hebbachi, Marina Kalinina, Annika Hausharter, Sébastien Favre, Simon Lebaron, Sherif Ismail, Timo Denk, Katharina Schlick, Thomas Fröhlich, Priya Bhutada, Erik Farquar, Katharina Schindlmaier, Sara Sormaz, Ed Hurt, Dieter Kressler, Yves Henry, Sarah A. Woodson, François Dragon, Roland Beckmann, Anthony K. Henras, Brigitte Pertschy

## Abstract

The early steps of eukaryotic large ribosomal subunit assembly remain poorly understood due to the structural flexibility of pre-60S intermediates, whose rRNA is extensively modified by small nucleolar RNPs (snoRNPs). Some snoRNPs, however, lack any modification function and instead scaffold ribosome assembly through largely unknown mechanisms. Here, we show that the H/ACA snoRNP snR37 integrates both modifying and scaffolding roles. Biochemical and structural analyses reveal a canonical H/ACA core that pseudouridylates a conserved uridine in the A site of the peptidyl transferase center, the catalytic heart of the 60S subunit. Additional RNA helices recruit non-core proteins, the Upa1-Upa2 heterodimer and Rbp95, which mediate stable snR37 association with pre-60S complexes. These proteins cooperate with the Npa1 rRNA chaperone complex to link four rRNA domains, thereby structurally organizing early pre-60S intermediates and promoting proper formation of the PTC. This dual organization establishes a paradigm for snoRNPs combining rRNA modification and scaffolding functions.

## Introduction

Ribosome biogenesis is one of the most energy-intensive and intricately coordinated processes in eukaryotic cells. It is highly conserved and has been most comprehensively characterized in the budding yeast *Saccharomyces cerevisiae*. Ribosome biogenesis begins in the nucleolus with the synthesis of a long ribosomal RNA precursor (pre-rRNA) by RNA polymerase I. This pre-rRNA contains the 18S, 5.8S, and 25S rRNAs, which are separated and flanked by spacer sequences that are sequentially removed during maturation. Assembly factors (AFs) and ribosomal proteins associate co-transcriptionally with the nascent transcript to form the earliest precursor particle, termed 90S particle or SSU processome. Subsequently, rRNA cleavages and extensive remodeling events give rise to precursors of the small (40S) and large (60S) ribosomal subunits. These particles follow distinct maturation pathways as they transit from the nucleolus to the nucleoplasm and ultimately to the cytoplasm, where final maturation steps yield functional ribosomal subunits (reviewed in ^1–6^). While intermediate and late maturation events have been described in considerable detail, the molecular mechanisms that orchestrate early pre-rRNA folding and domain compaction remain incompletely understood.

During these early maturation stages, when pre-ribosomal particles are still structurally flexible, numerous small nucleolar ribonucleoprotein particles (snoRNPs) associate with the pre-rRNA. snoRNPs belong to two major families, box C/D and box H/ACA, each consisting of a guide snoRNA associated with a set of core proteins. They have traditionally been studied for their catalytic function in installing site-specific post-transcriptional modifications, 2′-*O*-ribose methylation and pseudouridylation, respectively. Notably however, three yeast snoRNPs, U3, snR30, and snR190, deviate from this paradigm by fulfilling essential structural roles in early ribosome assembly. U3 coordinates 90S particle assembly and prevents premature formation of the central pseudoknot within the 18S rRNA.^7–18^ snR30 mediates the independent assembly of the 18S rRNA platform domain, while snR190 is believed to facilitate rRNA folding in early pre-60S particles.^19–24^ These non-catalytic snoRNPs, characterized by additional specific protein components, have long been considered specialized exceptions.^22, 25–29^ Whether structural functions extend beyond this small subset has remained unclear.

Early pre-rRNA folding also depends on dedicated AFs. A central organizer of early pre-60S maturation is the Npa1 complex, whose precise molecular function remains elusive. The complex contacts three distinct domains (I, II and VI) of the 25S rRNA, associates with the snoRNA snR190, and contains the RNA helicase Dbp6.^23,28, 30–33^ In addition, it likely interacts with two additional RNA helicases, Dbp7 and Dbp9.^34, 35^ Collectively, these observations strongly support a model in which the Npa1 complex functions as an rRNA chaperone to coordinate early pre-60S folding.^23, 28, 31^ We recently identified the early pre-60S AF Rbp95 as a member of the functional network of the Npa1 complex.^36^

Here, we uncover two additional members of this network, Upa1 and Upa2. Together with Rbp95, they are integral components of the snR37 box H/ACA snoRNP. Strikingly, snR37 exhibits an unexpected dual function: in addition to guiding rRNA modification, it promotes early pre-60S assembly. Our findings suggest that assembly-promoting functions may be more widespread among modification snoRNPs than previously anticipated.

## Results

### Nucleolar proteins Upa1 and Upa2 form a heterodimer that binds to Rbp95

Rbp95 was recently identified as an early pre-60S AF that binds to helix H95 in domain VI of 25S rRNA and that is genetically linked to the Npa1 complex.^36^ It contains a predicted α-helical core flanked by unstructured N-and C-terminal extensions (**Figure 1**A). Affinity purification of Rbp95 led to strong co-enrichment of Npa1 complex members as well as two additional AFs, Upa1 and Upa2.^36, 37^ Upa1 and Upa2 are proposed paralogs with 70% sequence identity (**Figure S1**A), sharing an OB (oligonucleotide/oligosaccharide-binding) fold and domains specific for SPOUT methyltransferases (**Figure 1A**).^38^ Consistent with large-scale localization data^39^, GFP-tagged Upa1 and Upa2 co-localized with the nucleolar marker Nop58, indicating their steady-state cellular localization is in the nucleolus, where early stages of ribosome assembly take place (**Figure 1B**). Moreover, Upa1-TAP or Upa2-TAP co-purified pre-rRNAs characteristic of 90S (35S pre-rRNA) and early pre-60S (27SA_2_ and 27SB pre-rRNAs) particles (**Figure S1**C), in line with a previous study suggesting their association with early pre-60S particles.^37^ Previous work, as well as reciprocal pull-down experiments of Upa1-Myc and Upa2-Flag further supported that the two proteins occur together in the same complex (**Figure S1**D).^37^

**Figure 1.**
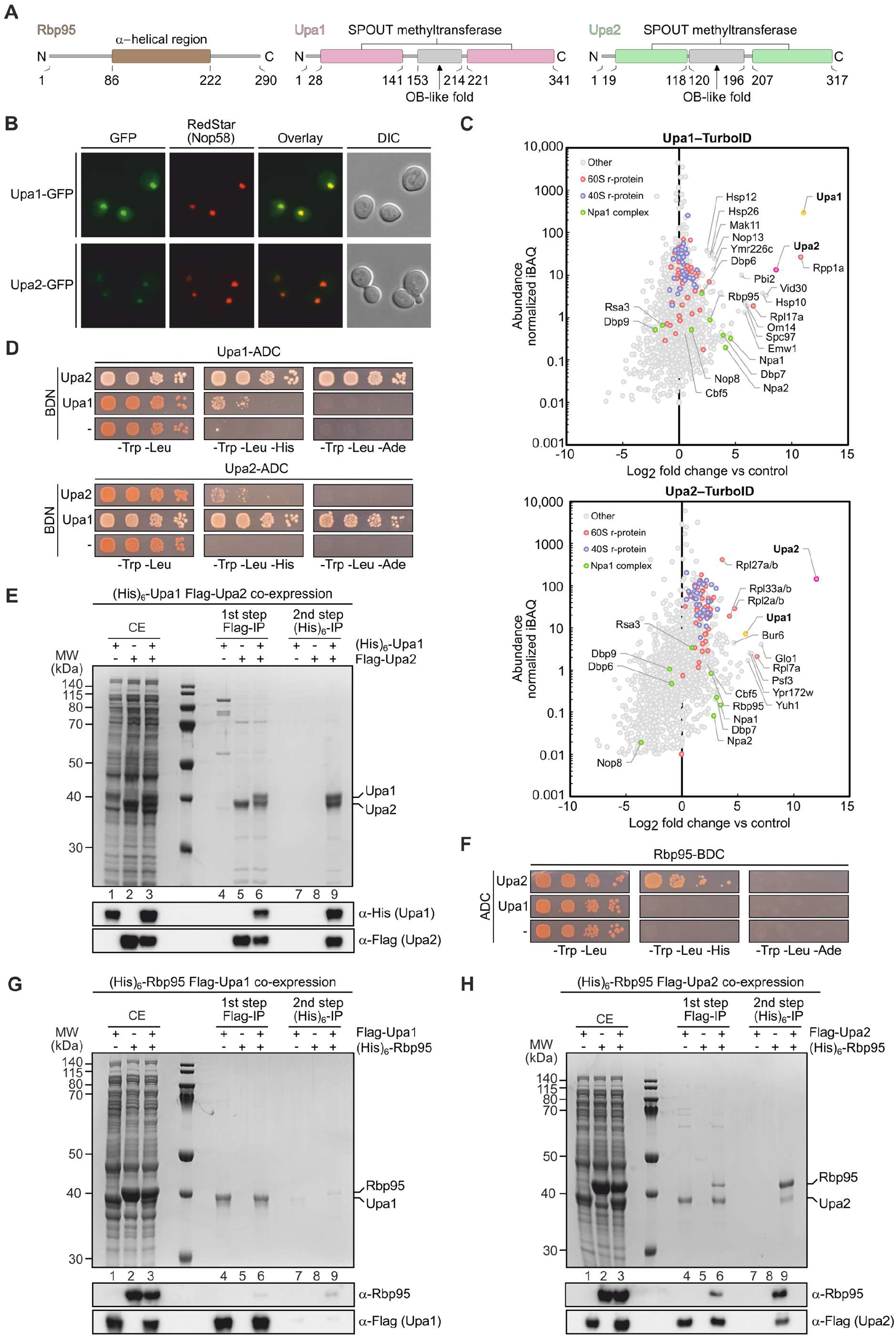
Upa1 and Upa2 are early ribosome biogenesis factors that form a heterodimer and physically interact with Rbp95. (**A**) Predicted domain organization of Rbp95, Upa1, and Upa2. (**B**) Fluorescence microscopy showing nucleolar localization of Upa1-GFP and Upa2-GFP. Nop58-RedStar was used as a nucleolar marker protein. DIC, differential interference contrast. Images were processed equally; an enhanced version of the Upa2-GFP image is shown in **Figure S1**B. (**C**) TurboID-based proximity labeling assays with the Upa1-TurboID and Upa2-TurboID baits. Detected proteins are plotted by normalized abundance (iBAQ, intensity-based absolute quantification; y-axis) and relative enrichment (log_2_-transformed fold change; x-axis) compared to a control (SV40NLS-yEGFP-TurboID). Enriched proteins appear on the right of the plot. (**D, F**) Y2H assays between Upa1, Upa2, and Rbp95. Bait proteins were either N-or C-terminally (BDN or BDC) fused to the Gal4 DNA-binding domain, and prey proteins were C-terminally fused to the Gal4 activation domain (ADC). Combinations with an empty vector (-) served as negative controls. Cells were spotted in serial dilution on SDC-Trp-Leu (growth control), SDC-Trp-Leu-His (indicating weak interaction), and SDC-Trp-Leu-Ade (indicating strong interaction). See **Figure S1**F for additional controls. (**E**) Interaction of (His)_6_-Upa1 and Flag-Upa2 co-expressed in *E. coli* and purified via sequential Flag and His affinity steps. Proteins expressed individually served as negative controls. Eluates (Flag-IP and His-IP) and crude extracts (CE) were analyzed by SDS-PAGE and Coomassie staining or western blotting using the indicated antibodies. (**G, H**) Upa2 directly interacts with Rbp95. Flag-Upa2 (G) or Flag-Upa1 (H) were co-expressed with (His)_6_-Rbp95 in *E. coli* and purified via sequential Flag and His pull-downs. Eluates (Flag-IP and His-IP) and CE were analyzed as in (E).

To identify proteins in physical proximity of Upa1 and Upa2 within pre-ribosomes, we employed TurboID-based proximity labeling.^40–43^ Notably, Upa2 was among the most prominent co-enriched proteins in the Upa1 TurboID experiments (irrespective of C-terminal or N-terminal positioning of the TurboID tag) (**Figures 1**C and **S1**E). The converse was also true, with Upa1 being co-enriched in the TurboID assays performed with C-or N-terminally TurboID-tagged Upa2. These results suggested that Upa1 and Upa2 likely interact directly. In addition, several early pre-60S AFs were enriched, including Rbp95 and members of the Npa1 complex, such as Npa1 and Npa2, and the Npa1 complex-associated RNA helicase Dbp7 (**Figure 1**C). These data suggested that Upa1 and Upa2 bind to pre-60S particles near the Npa1 complex and Rbp95.

The co-enrichment of Upa1 with Upa2 suggests that they are likely not functionally redundant paralogs, but instead act together in the same complex, in agreement with the usually dimeric nature of SPOUT proteins^44, 45^. Yeast two-hybrid (Y2H) assays indeed revealed that Upa1 and Upa2 strongly interact with each other (**Figure 1**D). Furthermore, co-expression of (His)_6_-Upa1 and Flag-Upa2 in *Escherichia coli* followed by sequential affinity purification confirmed that the two proteins form a stoichiometric complex (**Figure 1**E). Next, we investigated whether either protein directly physically interacts with Rbp95. Both Y2H and co-purification assays revealed an interaction of Rbp95 with Upa2, but not with Upa1 (**Figures 1**F-**1**H).

Together, these results establish pre-60S AFs Upa1 and Upa2 as a heterodimer, with Upa2 additionally mediating the interaction with Rbp95.

### Upa1 and Upa2 are associated with the snR37 H/ACA snoRNP

To reveal whether Upa1 and Upa2 are associated with early pre-60S particles by directly interacting with rRNA, we performed RNA crosslinking and analysis of cDNA (CRAC).^46^ Strikingly, the majority of reads obtained in the Upa1-CRAC experiment (∼80%) and a substantial proportion of reads obtained for the Upa2-CRAC experiments (∼20%) mapped to a single H/ACA snoRNA, snR37, indicating a stable and specific association of both proteins with the snR37 snoRNP (**Figure 2**A). Importantly, we previously detected crosslinking of Rbp95 to several snoRNAs, with the highest number of snoRNA reads originating from snR37.^36^ Together with the physical interaction between Rbp95 and Upa2 (**Figures 1**F and **1**H), these results suggested that Rbp95 is also part of the snR37 snoRNP interaction network. The association of snR37 with specific proteins besides the H/ACA core components classifies it as an unconventional snoRNP and identifies it as a strong candidate for harboring functions that extend beyond rRNA modification.

**Figure 2.**
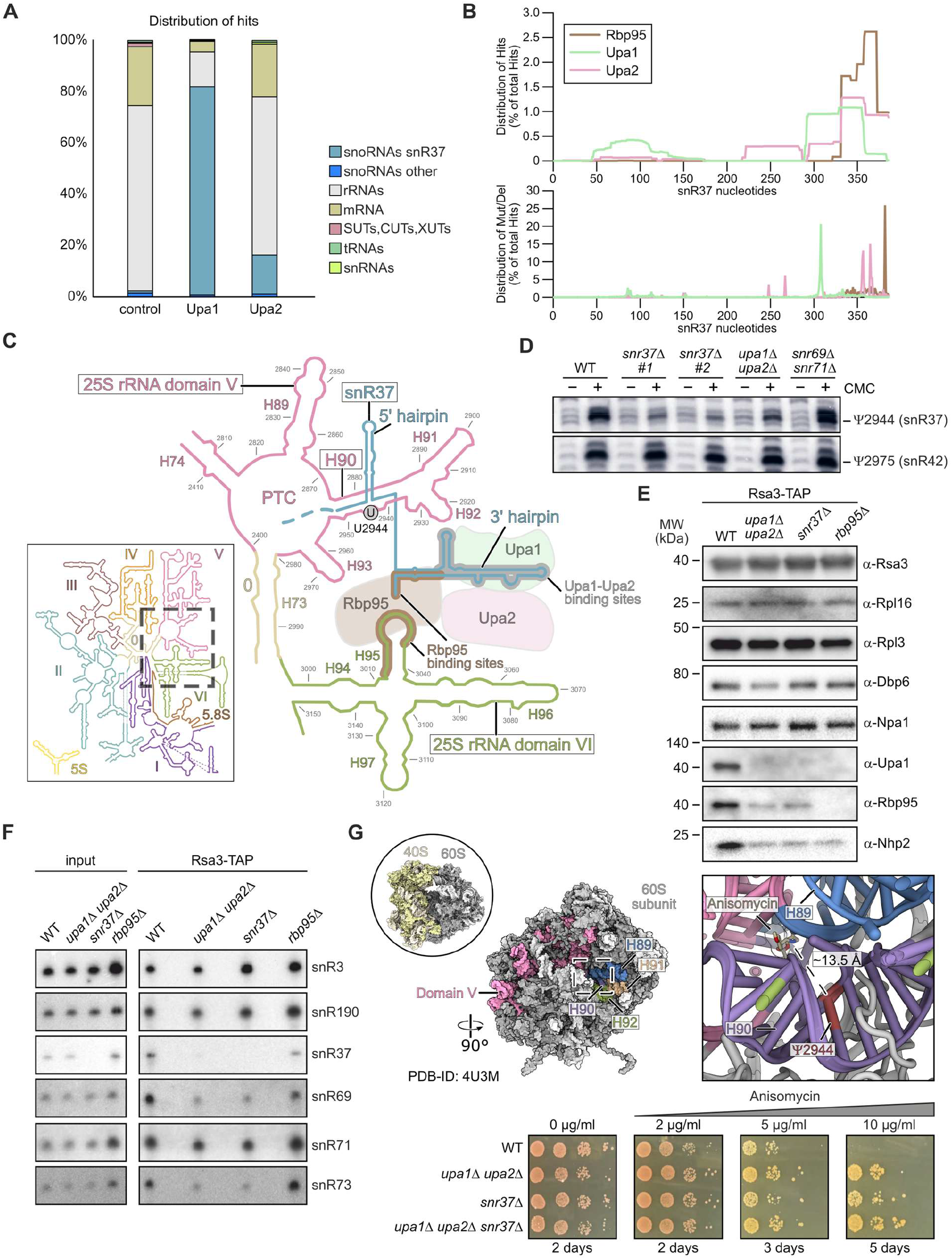
Upa1 and Upa2 interact with the H/ACA snoRNA snR37. (**A**) CRAC analyses of Upa1-HTP and Upa2-HTP reveal binding to snR37. UV-crosslinked RNA-protein complexes were isolated, RNAs reverse transcribed, and sequenced. A wild-type strain (BY4741) served as control. (**B**) Upper panel: Representation of each nucleotide of snR37 in the sequences obtained in the Upa1, Upa2, and Rbp95 CRAC experiments. Lower panel: Representation of mutations/deletions in the sequences, marking likely actual crosslinking sites. (**C**) Schematic representation of snR37 snoRNP interactions with the 25S rRNA and with Rbp95, Upa1, and Upa2. The depicted region of the 25S rRNA is highlighted within the box showing the overall 25S rRNA secondary structure. snR37 base-pairs through its 5′ hairpin with helix H90 in domain V of the 25S rRNA to guide pseudouridylation at U2944. In addition, snR37 associates with helix H95 in domain VI via Rbp95. Rbp95, Upa1, and Upa2 bind to the 3′ hairpin of snR37. (**D**) Primer extensions after CMC treatment of RNA from wild-type (WT) and mutant strains *upa1*Δ *upa2*Δ, *snr37*Δ (two clones), and *snr69*Δ *snr71*Δ. Ψs cause reverse transcription stops in CMC-treated samples. Full gel: **Figure S3**A. (**E**) Western blot analysis of Rsa3-TAP purification eluates from wild-type (WT) and deletion strains *upa1*Δ *upa2*Δ, *snr37*Δ, and *rbp95*Δ. Inputs are shown in **Figure S3**B. (**F**) Northern blot analysis of RNAs extracted from Rsa3-TAP purifications in (E) using probes against selected snoRNAs. snR3, which binds to a different 25S rRNA domain (domain IV) and should therefore be unaffected, was used as loading control. Quantifications of the result are shown in **Figure S3**C. (**G**) Upper panels: Surface view of the 80S ribosome bound to anisomycin (PDB-ID: 4URM) and the corresponding 60S subunit, highlighting domain V including PTC helices H89-H92. Close-up shows the molecular model of the PTC including anisomycin and the position of Ψ2944. The distance between anisomycin and Ψ2944 is indicated with a dashed line. Lower panels: Growth assay of wild-type (WT) and mutant cells (*upa1*Δ *upa2*Δ, *snr37*Δ, or *upa1*Δ *upa2*Δ *snr37*Δ) on YPD plates with increasing anisomycin concentrations (0, 2, 5, or 10 µg/ml). Plates were incubated at 30 °C for 2 (0 and 2 µg/ml), 3 (5 µg/ml) or 5 (10 µg/ml) days.

Mapping of the RNA sequences crosslinked to Upa1, Upa2, and Rbp95 (**Figure 2**B, upper panel), together with the positions of mutations in the reads, which indicate the precise crosslinking sites (**Figure 2**B, lower panel) revealed that all three proteins bind primarily to the 3′ region of snR37. Sequences corresponding to snR37 nucleotides 50 to 150 were also enriched in the Upa1 CRAC experiment, but to a lower extent.

Notably, although Rbp95 also crosslinked with snR37, its primary crosslinking site mapped to helix H95 in domain VI of the 25S rRNA.^36^ This site is located less than 100 nucleotides downstream of the sole residue predicted to be pseudouridylated by the snR37 snoRNP, U2944, which resides in helix H90 within 25S rRNA domain V.^47, 48^ Base pairing of snR37 with helix H90, combined with Rbp95 binding to helix H95, would allow to tether domains V and VI of the 25S rRNA (**Figures 2**C and **S2**). To test whether snR37 indeed directs pseudouridylation at U2944, we performed primer extension analysis following CMC (N-Cyclohexyl-N′-(2-morpholinoethyl)carbodiimide) treatment (**Figures 2**D and **S3**A). The primer extension signal arising from Ψ2944 was abrogated in a *snr37*Δ strain, confirming that snR37 guides pseudouridylation at this position. The signal was also reduced in a *upa1*Δ *upa2*Δ double mutant, indicating that Upa1 and Upa2 are required for snR37-guided modification. In contrast, pseudouridylation at other positions, such as U2975, which is guided by snR42 H/ACA snoRNA, remained unaffected (**Figures 2**D and **S3**A). Furthermore, the absence of snR69 and snR71, two box C/D snoRNAs whose binding sites on helix H90 overlap with that of snR37 (**Figure S2**), had no effect on the CMC-dependent primer extension signal at U2944 (**Figure 2**D). These results demonstrate that the snR37 snoRNP is responsible for pseudouridylation at U2944, and that this function depends on Upa1 and Upa2.

### Upa1, Upa2, and Rbp95 are required for stable pre-60S association of snR37

The requirement of Upa1 and Upa2 for pseudouridylation at U2944 could reflect either a direct role in modification or an indirect role, for example by promoting stable association of snR37 with pre-60S particles. To test whether snR37, Upa1, Upa2, and Rbp95 depend on each other for pre-60S binding, we analyzed the protein (**Figure 2**E) and snoRNA (**Figure 2**F) composition of early pre-60S particles purified from *snr37*Δ, *upa1*Δ *upa2*Δ, and *rbp95*Δ mutants. Upa1 levels were drastically reduced in pre-60S particles isolated from the *snr37*Δ strain, consistent with previous results,^37^ and were similarly also reduced in pre-60S particles purified from the *rbp95*Δ strain (**Figures 2**E). Deletion of either *SNR37* or both *UPA1* and *UPA2* resulted in reduced association of Rbp95 with pre-60S particles. Also, the levels of the H/ACA snoRNP core component Nhp2 were reduced in all three mutants. Analysis of snR37 levels in early pre-60S particles showed that, although snR37 was stable in the *upa1*Δ *upa2*Δ strain (**Figure 2**F, input), it was absent from the purified pre-60S particles (**Figure 2**F and **S3**C). A clear reduction of snR37 was also observed in pre-60S particles purified from a *rbp95*Δ strain. Together, these results indicate that snR37, Upa1-Upa2, and Rbp95 depend on each other for efficient association with pre-60S particles, consistent with their functioning as a pre-assembled complex.

Several box C/D snoRNAs targeting the same region of helix H90 were also affected. In particular, the levels of snR69, snR71, and, most prominently, snR73, whose binding sites overlap with (snR69 and snR71) or lie adjacent to (snR73) the snR37 base-pairing region, were reduced in pre-60S particles from *snr37*Δ and *upa1*Δ *upa2*Δ strains, and to a lesser extent from the *rbp95*Δ strain. In contrast, the nearby snoRNA snR190 was unaffected (**Figures 2**F, **S2**, and **S3**C). Consequently, the modifications guided by snR69 and snR73 (2′-*O*-ribose methylations at C2948 and C2959), and to a lesser extent the modification guided by snR71 (2′-*O*-ribose methylation at A2946), were reduced in *snr37*Δ and *upa1*Δ *upa2*Δ mutants (**Figure S3**D). These findings suggest that loss of snR37 may induce local structural changes in helix H90 that impair binding of other snoRNAs targeting this region.

U2944, the modification site of snR37 in helix H90, is part of the A site of the peptidyl transferase center (PTC) that positions the aminoacyl-tRNA for peptide bond formation during translation. Mutants lacking snoRNAs that target the PTC have previously been reported to show altered sensitivity to the A-site inhibitor anisomycin.^49–51^ Consistent with this, both the *snr37*Δ mutant and the *upa1*Δ *upa2*Δ double mutant exhibited anisomycin resistance (**Figure 2**G), potentially reflecting structural alterations in the A site. Additionally, *upa1*Δ *upa2*Δ cells showed a mild translation defect, as indicated by their reduced polysome content compared to wild-type cells (**Figure S3**E). Collectively, our results suggest that loss of the snR37 snoRNP induces structural changes in the PTC, conferring resistance to anisomycin and leading to reduced association of other snoRNAs that normally bind in this region.

### Upa1, Upa2, and snR37 cooperate with the Npa1 complex

We next investigated whether the snR37 snoRNP also interacts with proteins within the pre-60S particle, particularly through its associated protein components Upa1 and Upa2. As our TurboID proximity labeling experiments suggested that Upa1 and Upa2 are close to the Npa1 complex on pre-60S particles (**Figure 1**C), we examined potential physical interactions between Upa1 or Upa2 and components of the Npa1 complex by Y2H assays (**Figures 3**A and **S4**). These experiments revealed Y2H interactions between Upa1 and the Npa1 complex members Npa2 and Dbp6, whereas no interactions were detected for Upa2 with any of the tested proteins. Altogether, our protein-protein interaction data (**Figures 1**D-**1**H and **3**A) suggest that within the Upa1-Upa2 heterodimer, Upa2 interacts with Rbp95, while Upa1 mediates contact with the Npa1 complex, particularly with Dbp6 and Npa2 (**Figure 3**B). Base-pairing of snR37 to 25S rRNA domain V and binding of its interactor Rbp95 to domain VI would connect these two domains, whereas the Npa1 complex is known to tether domains I, II, and VI. Hence, the physical interaction between snR37 snoRNP-associated proteins and the Npa1 complex could facilitate the spatial clustering of domains I, II, V, and VI, promoting compaction of the 25S rRNA during early stages of 60S assembly.

**Figure 3.**
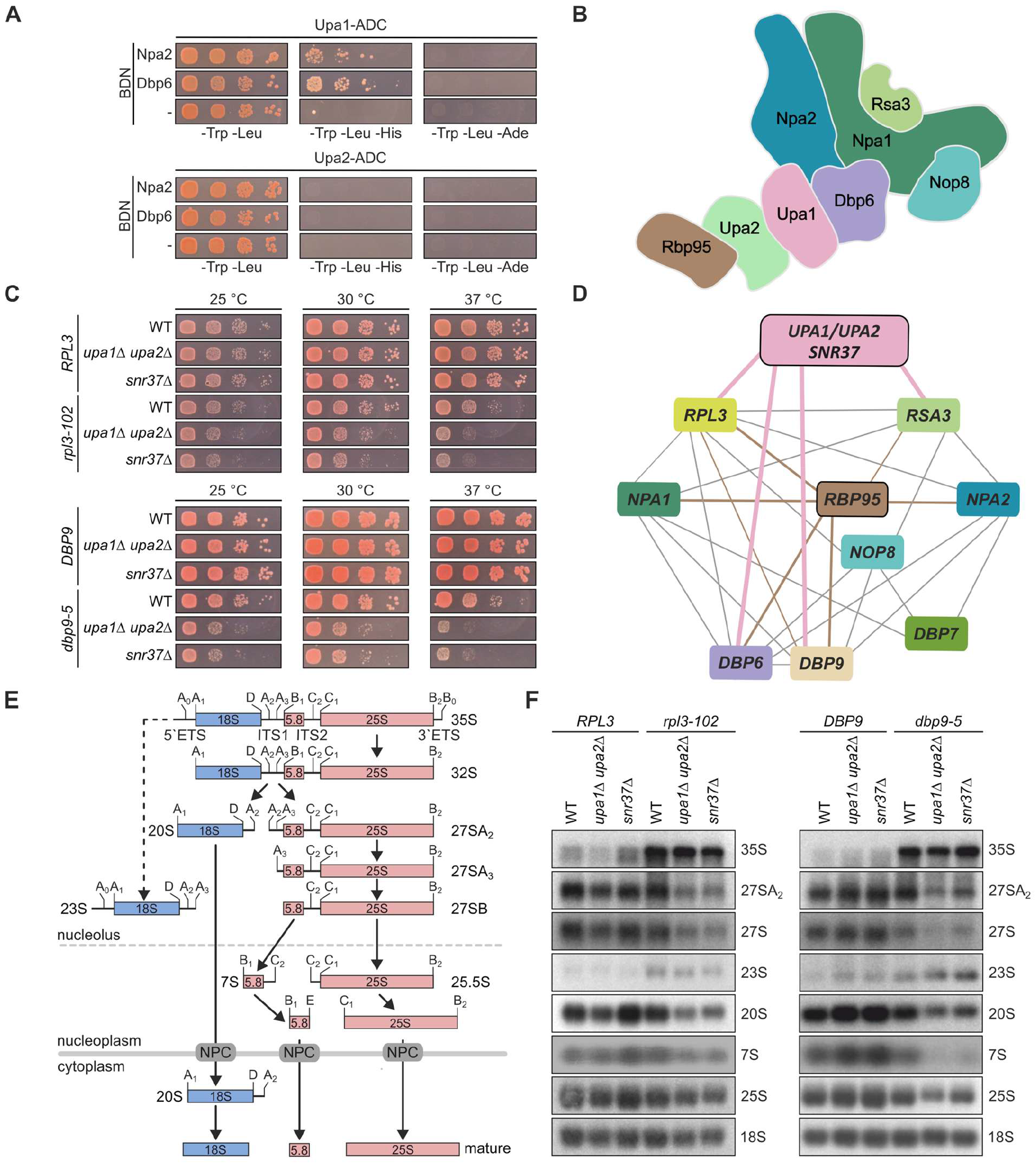
Upa1, Upa2, and snR37 are physically and genetically linked to the Npa1 complex. (**A**) Y2H assays between Upa1 or Upa2, C-terminally fused to the Gal4 activation domain (ADC), and Npa1 complex members Npa2 and Dbp6, fused N-terminally to the Gal4 DNA-binding domain (BDN). Combinations with an empty vector (-) served as negative controls. Cells were spotted in serial dilution on SDC-Trp-Leu (growth control), SDC-Trp-Leu-His (indicating weak interaction), and SDC-Trp-Leu-Ade (indicating strong interaction). See **Figure S4** for additional controls. (**B**) Model of physical interactions: Upa1 and Upa2 form a heterodimer connecting Rbp95 with the Npa1 complex. (**C**) Deletion of *UPA1*/*UPA2* or *SNR37* exacerbates growth defects of *rpl3* and *dbp9* mutants. Yeast cells (*rpl3*Δ or *dbp9*Δ) carrying *LEU2* plasmids with either wild-type or mutated alleles of *RPL3* or *DBP9* in combination with *UPA1* and *UPA2* or *SNR37* deletions were spotted on SDC-Leu plates and incubated for 3 days at the indicated temperatures. (**D**) *UPA1*-*UPA2* and *SNR37* are part of the *NPA1* genetic network. Genetic interactions identified here (**Figures 3**C **S5**D, and **S5**E; pink), previously identified *RBP95* genetic interactions (brown)^36^, and additional known *NPA1* network links (gray)^89,33, 90, 34, 91, 30^ are indicated. (**E**) Schematic overview of rRNA processing steps during ribosome biogenesis. (**F**) Northern blot analysis of total RNA extracted from yeast cells with the indicated genotypes from (C). rRNA processing defects were detected by northern blotting using probes for 25S, 18S, A2-A3 (27SA_2_, 35S, and 23S pre-rRNAs), E-C2 (27SA, 27SB, and 7S pre-rRNAs), and D-A2 (20S pre-rRNA).

We observed that cells individually or simultaneously lacking Upa1 and Upa2 in the presence or absence of snR37 or Rbp95 did not exhibit any noticeable growth defect (**Figures S5**A-**S5**C). Given the physical link to the Npa1 complex, we hypothesized that functional redundancies between snR37 snoRNP and the Npa1 complex might buffer phenotypes arising from the loss of snR37 or its associated components. Since *RBP95* was previously shown to genetically interact with members of the *NPA1* genetic network,^36^ we extended this analysis to test whether *UPA1, UPA2*, or *SNR37* also show genetic interactions with members of this network-*RPL3* (coding for uL3), *DBP9, DBP6*, or *RSA3* (**Figures 3**C, **S5**D, and **S5**E). Indeed, deletion of *UPA1* and *UPA2*, or *SNR37* enhanced the mild growth defects of *rpl3-102* and *dbp9-5* mutants (**Figure 3**C). Additional genetic interactions were observed between *upa1*Δ or *upa2*Δ and *dbp6-2* and *dbp6-3* mutants (**Figure S5**D), as well as between the *upa1*Δ *upa2*Δ double mutant and *rsa3*Δ (**Figure S5**E). To conclude, our data suggest that *RBP95, UPA1, UPA2*, and *SNR37* are all part of the *NPA1* genetic network (**Figure 3**D), further supporting partial functional redundancy between the snR37 snoRNP and the Npa1 complex.

If the snR37 snoRNP indeed acts in concert with the Npa1 complex during early pre-60S maturation, the observed synthetic growth defects should coincide with enhanced ribosome biogenesis defects. To test this, we monitored pre-rRNA processing intermediates (**Figures 3**E and **3**F). Deletion of *UPA1* and *UPA2* or *SNR37* alone did not cause any detectable pre-rRNA processing defects. The *rpl3-102* and *dbp9-5* mutants accumulated 35S pre-rRNA, and *dbp9-5* additionally showed reduced overall 27S levels (corresponding to the 27SA_2_, 27SA_3_, and 27SB pre-rRNA species) and, to a lesser extent, 20S pre-rRNA levels. When combined with deletion of *UPA1* and *UPA2* or *SNR37*, these mutants exhibited a pronounced reduction of the 27SA_2_ pre-rRNA, indicating reduced synthesis or stability of the earliest pre-60S precursors. Because 27SA_2_ and 20S pre-rRNA are generated simultaneously from cleavage of the 32S pre-rRNA, the decrease in 20S pre-rRNA levels further supports defects in early processing steps. These early defects propagated to later stages of ribosome assembly, as indicated by reduced levels of downstream pre-rRNAs (see total 27S signal and 7S pre-rRNAs) and a marked decrease in mature 25S rRNA relative to 18S rRNA, pointing to a specific impairment of 60S subunit maturation (**Figure 3**F).

Taken together, our findings show that the snR37 snoRNP cooperates with the Npa1 complex during early stages of pre-60S maturation.

### Architecture of the snR37 snoRNP

To establish whether Upa1, Upa2, and Rbp95 function as adapter proteins for the recruitment of the snR37 snoRNP to early pre-60S particles or instead represent integral protein components of the snR37 snoRNP itself, we performed affinity purification of genomically expressed Upa1 bearing a C-terminal FTpA tag, followed by sucrose gradient sedimentation analysis (**Figure 4**A). This approach revealed two distinct populations: early pre-60S particles sedimenting in the heavier fractions (9-11), and a smaller, less complex particle in the lighter fractions (4 and 5).

**Figure 4.**
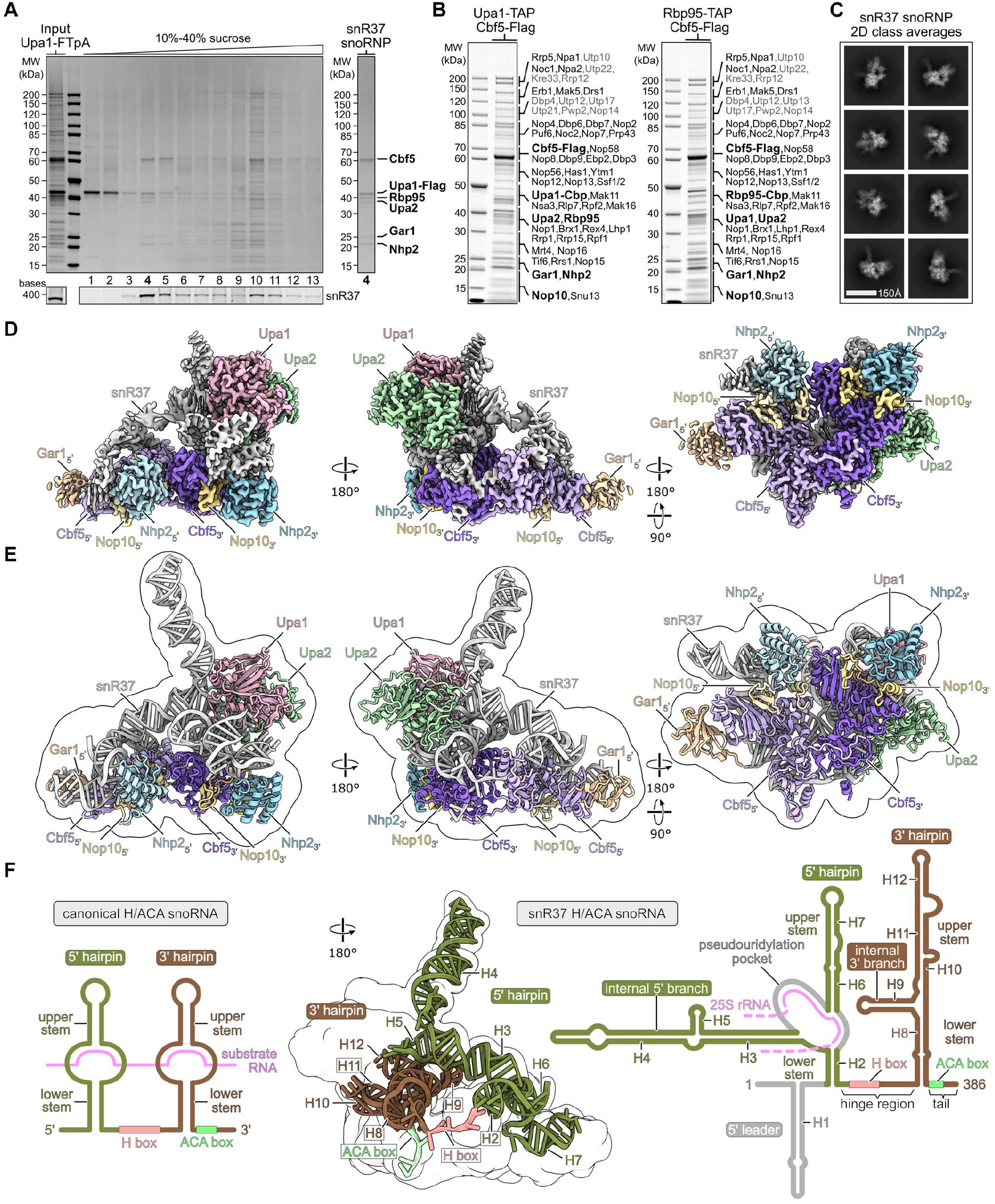
Structure of the unconventional snR37 H/ACA snoRNP. (**A**) Sucrose gradient analysis of Upa1-FTpA affinity-purified particles. The Coomassie-stained SDS-PAGE shows free snoRNPs in fractions 4-5 and pre-60S particles associated with Upa1 in fractions 9-11. The input is shown on the left. SYBR green was used to detect snR37. The free snoRNP in lane 4 is also shown on the right and contains H/ACA core proteins, Upa1, Upa2, and Rbp95. (**B**) Coomassie-stained SDS-PAGE of split-tag affinity purifications of complexes purified via Upa1-Cbf5 and Rbp95-Cbf5. Proteins identified by mass spectrometry are indicated; pre-60S AFs are written in black, 90S factors in gray. Components of the snR37 snoRNP are highlighted in bold with increased font size. (**C**) 2D class averages of the snR37 snoRNP (scale bar 150 Å). (**D, E**) Cryo-EM density map (D) and molecular model (E) of the free snR37 snoRNP in different orientations. Associated proteins are colored and labeled. Silhouette of the filtered cryo-EM map is overlayed with the molecular model (E). (**F**) Secondary structure comparison between a canonical H/ACA snoRNA (left) and the snR37 H/ACA snoRNA (right). The snR37 snoRNP structure is shown with the snoRNA highlighted as molecular model and the protein factors shown as filtered, transparent surface (middle). The RNA elements of the snR37 snoRNA, including the 5′ and 3′ hairpins, H-and ACA boxes, and RNA helices (H1-H12), are labeled. Regions of the snR37 snoRNA that could not be modeled are shown in gray.

snR37 snoRNA was enriched in both populations. Subsequent mass spectrometry analysis of the proteins within the smaller complex (fraction 4) identified the canonical H/ACA core proteins Cbf5, Gar1, and Nhp2. The fourth core component, Nop10, was not detected, likely due to its small size (∼7 kDa), which may cause it to migrate out of the gel during electrophoresis. Upa1, Upa2, and Rbp95 were also present in the complex, indicating that they are integral non-core components of the free snR37 snoRNP (**Figure 4**A, right panel).

To isolate the snR37 snoRNP for structural analysis by cryo-EM, we employed a split-tag affinity purification strategy. Upa1 or Rbp95 fused to a TAP tag were used as baits in the first step, followed by purification via the H/ACA core protein Cbf5 fused to a Flag tag. This approach allowed us to enrich both free and pre-60S bound forms of the snR37 snoRNP (**Figure 4**B). Cryo-EM analysis of these preparations revealed distinct ∼170 Å particles corresponding to the free snoRNPs, while no useful 2D class averages of early pre-60S intermediates could be obtained, likely due to their intrinsic flexibility. 2D class averages of the free snoRNPs revealed globular particles with clear features for the two H/ACA protomers as well as a helical thread-like extension (**Figure 4**C). Individual datasets produced highly similar classes and were therefore combined to increase particle numbers. Non-uniform refinement and 3D classification revealed several closely related conformations with slight changes in the relative orientation of 5′ and 3′ hairpin protomers and flexibility of the 5′ binding Gar1 (**Figure S6**). The region surrounding the pseudouridylation pocket of the 5′ hairpin was highly heterogeneous. Nevertheless, we isolated a stable class that was reconstructed at ∼2.8 Å resolution, allowing us to build a structural model of the snR37 snoRNP (**Figures 4**D,**4**E, **S6, S7**, and **Table S3**).

Canonical H/ACA snoRNAs consist of two hairpins (**Figure 4**F, left) that each contain a lower and an upper stem separated by a single-stranded pseudouridylation pocket that binds and positions the substrate RNA. The 5′ hairpin is followed by a hinge region containing the H box (ANANNA), while the 3′ hairpin is followed by the ACA box. With a length of 386 nucleotides, the snR37 snoRNA is much longer than canonical H/ACA snoRNAs (∼150-200 nucleotides) and includes several additional RNA elements (**Figures 4**F, right, and **S8**). The 5′ hairpin of snR37 is unusually long (204 nucleotides) and contains a long internal branch (internal 5′ branch) emerging from the unpaired region comprising the pseudouridylation pocket. In contrast, the 3′ hairpin lacks the unpaired bulge that would typically correspond to its pseudouridylation pocket. Instead, the lower stem is extended and the bulge is replaced by a three-helix junction (internal 3′ branch). Consequently, the architecture of the 3′ hairpin prevents substrate RNA binding in the enzymatic pocket of the 3′-binding Cbf5 copy (referred to as 3′ Cbf5). This leads to the delocalization of the Cbf5 thumb loop and sterically hinders the binding of the 3′-binding Gar1 copy (see below). A 5′ leader sequence also present in other snoRNAs, such as snR30,^21^ was not visible in the cryo-EM structure, likely due to conformational flexibility.

The snR37 snoRNP contains two H/ACA core protein modules built around a Cbf5 dimer through direct contact between the Cbf5 pseudouridine synthase and archaeosine transglycosylase (PUA) domains and the binding of the unstructured N-terminal extension (NTE) of the 5′ Cbf5 copy to the 3′ Cbf5 PUA domain. Together with the Cbf5 C-terminal extensions (CTEs), they interact with the H and ACA consensus motifs of snR37 and coordinate the lower stems of the 5′ and 3′ hairpins (**Figures 5**A-**5**C and **5**E middle). The catalytic domain of each Cbf5 copy binds to Nop10, which in turn interacts with Nhp2; together, the three proteins interact with and coordinate the upper stems of the snR37 hairpins (**Figures 5**D and **5**E). Overall, this arrangement resembles the typical H/ACA snoRNP core architecture as shown for the human telomerase H/ACA RNP, the structure of insect H/ACA snoRNPs, the yeast snR30 snoRNP as well as for canonical H/ACA RNPs from archaea (**Figure 5**E right).^22, 52–54^

**Figure 5.**
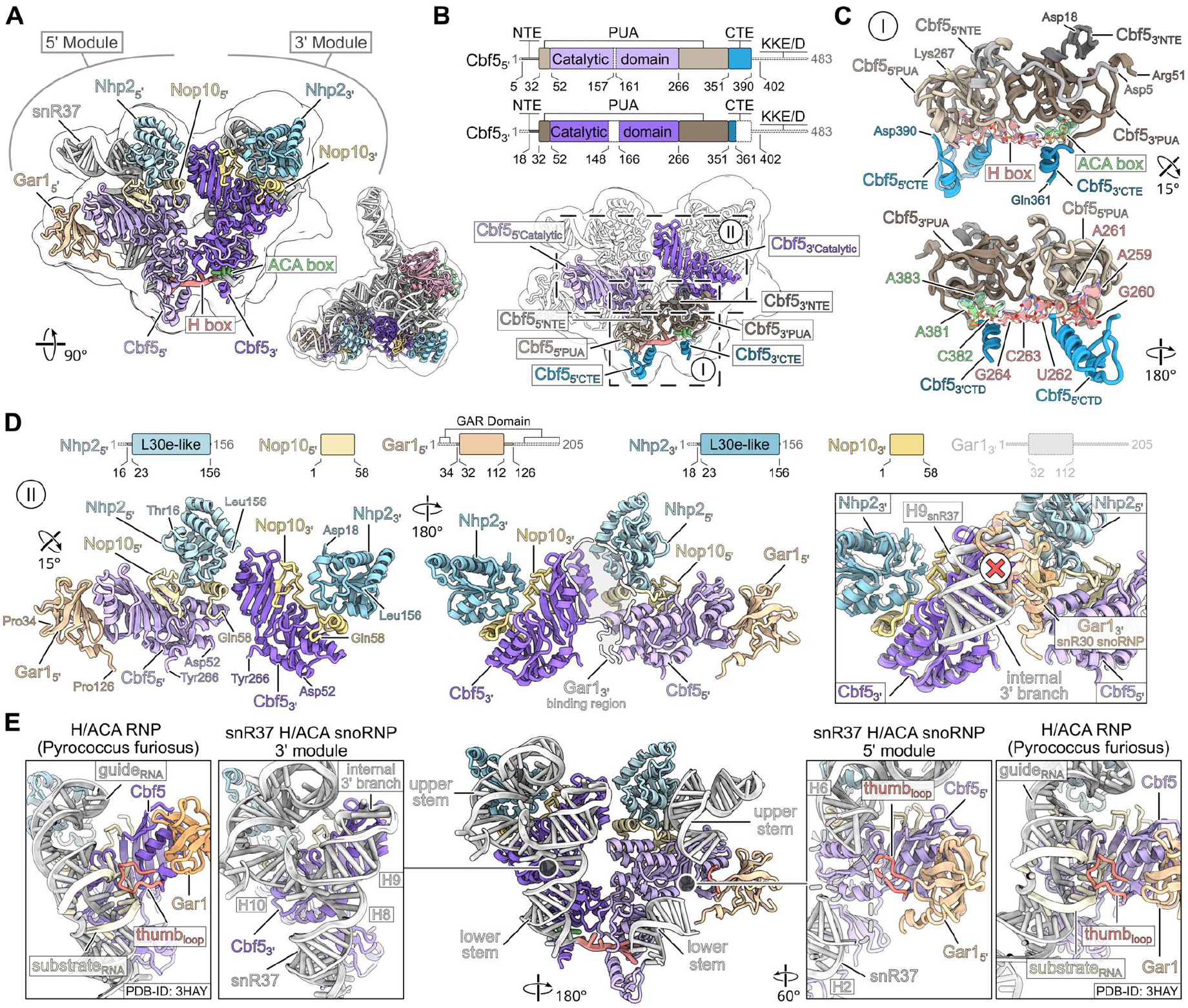
Structural analysis of the snR37 H/ACA core. (**A**) Molecular model and filtered density map of the H/ACA core. The 5′ and 3′ modules are indicated. (**B**) Domain architecture and molecular model of the Cbf5 dimer. The PUA and catalytic domain as well as the NTE and CTE (N-and C-terminal extensions) are indicated. Dashed lines indicate regions of Cbf5 missing in the molecular model, including the KKE/D-containing intrinsically disordered region (IDR). (**C**) Close-up of the Cbf5 PUA domain dimer showing its interactions with the H and ACA box consensus sequences of snR37, presented in two orientations. The H and ACA box models are overlaid with segmented, transparent cryo-EM densities. (**D**) The domain architectures of Nhp2, Nop10, and Gar1 are shown in the top panel. Regions of the proteins missing from the molecular model are shown as dashed lines, including the GAR domain IDRs of Gar1. The molecular models of the two Cbf5 catalytic domains and their interactions with Nop10-Nhp2 and Gar1 are shown in two orientations in the lower panel (left and middle). Magnified view illustrating the steric block of the Gar1-binding site within the 3′ module by the internal 3′ branch (H9) of snR37. The snR30 snoRNP structure (PDB-ID: 9G25) was rigid-body fitted and is shown as transparent gray molecular model. The 3′ Gar1 copy of the snR30 snoRNP clashing with the snR37 H9 of the snR30 snoRNP is shown in transparent orange (lower right panel). (**E**) Putative substrate-binding regions within the 3′ and 5′ H/ACA modules compared with the substrate RNA-bound H/ACA RNP from the archaeon *Pyrococcus furiosus* (PDB-ID: 3HAY). An overview is shown in the middle, with detailed views on the left and right. The archaeal model was fitted to either the 3′ or 5′ module, both of which are shown in the same orientation. The snR37 model of the internal 5′ hairpin (nt 62-174) has been removed to improve visualization.

The 5′ module, which guides pseudouridylation at U2944 in the 25S rRNA, contains the full complement of canonical core proteins (Cbf5, Gar1, Nop10, and Nhp2). Gar1’s globular domain (residues 32-112) binds to the catalytic domain of the 5′ Cbf5 in proximity to the active site where it is known to enhance the catalytic activity of Cbf5 and induce RNA substrate release (**Figure 5**D).^55, 56^ In our structure, the single-stranded regions of the pseudouridylation pocket are delocalized, likely because of the absence of substrate RNA (**Figure 5**E). Deviating from the canonical composition, the atypical 3′ protomer lacks the Gar1 subunit because its binding site is occupied by the extended lower stem (H8) and the internal 3′ branch helix (H9) of the snR37 3′ hairpin (**Figures 5**D and **5**E). Both snoRNA elements prevent substrate rRNA binding within the 3′ module, as evidenced by comparison of the snR37 snoRNP model with the X-ray structure of an archaeal H/ACA RNP bound to substrate RNA (**Figure 5**E, right).^52^

### Upa1-Upa2 interactions in the snR37 snoRNP

In line with our biochemical data and CRAC results, the snR37 snoRNP structure revealed that Upa1 and Upa2 are integral components of the atypical snR37 3′ module (**Figure 6**A). Both proteins belong to the SPOUT methyltransferase family and contain the characteristic α/β fold with a trefoil knot.^45^ Additionally, each protein contains an OB-like fold embedded within the SPOUT domain (**Figures 1**A and **6**B). Consistent with our *in vitro* data (**Figure 1**E) and like most SPOUT methyltransferases,^44, 45^ Upa1 and Upa2 form a dimeric structure. The Upa1-Upa2 complex has an antiparallel arrangement, with the OB-like folds placed on opposite sides of the complex. Hereby, two parallel α-helices within each of the SPOUT domains of Upa1 and Upa2 dimerize in an almost perpendicular arrangement (**Figure 6**B). The human homolog of Upa1-Upa2, SPOUT1 (CENP-32), forms a similar dimeric structurer (**Figures 6**C and **S9**A-**S9**C).^57^ The SPOUT1 homodimer has been crystallized in complex with the methyl donor S-adenosyl methionine (SAM) or its reaction product S-adenosyl-homocysteine (SAH), and functions as an active RNA methyltransferase.^57, 58^ However, comparison with SPOUT1 suggests that the Upa1-Upa2 heterodimer is enzymatically inactive when incorporated into the snR37 snoRNP (**Figures 6**C and **S9**A-**S9**C).^57^ In our structure, neither Upa1 nor Upa2 showed any density that could correspond to SAM. Moreover, sequence alignments show that residues important for formation of the SAM-binding pocket and for SAM interaction in SPOUT1 are not conserved in Upa1 and Upa2 (**Figure S9**A). Structural comparisons further reveal that Upa1-and Upa2-specific elements hinder the formation of the SAM-binding pocket within the snR37 snoRNP (**Figures S9**B and **S9**C).

**Figure 6.**
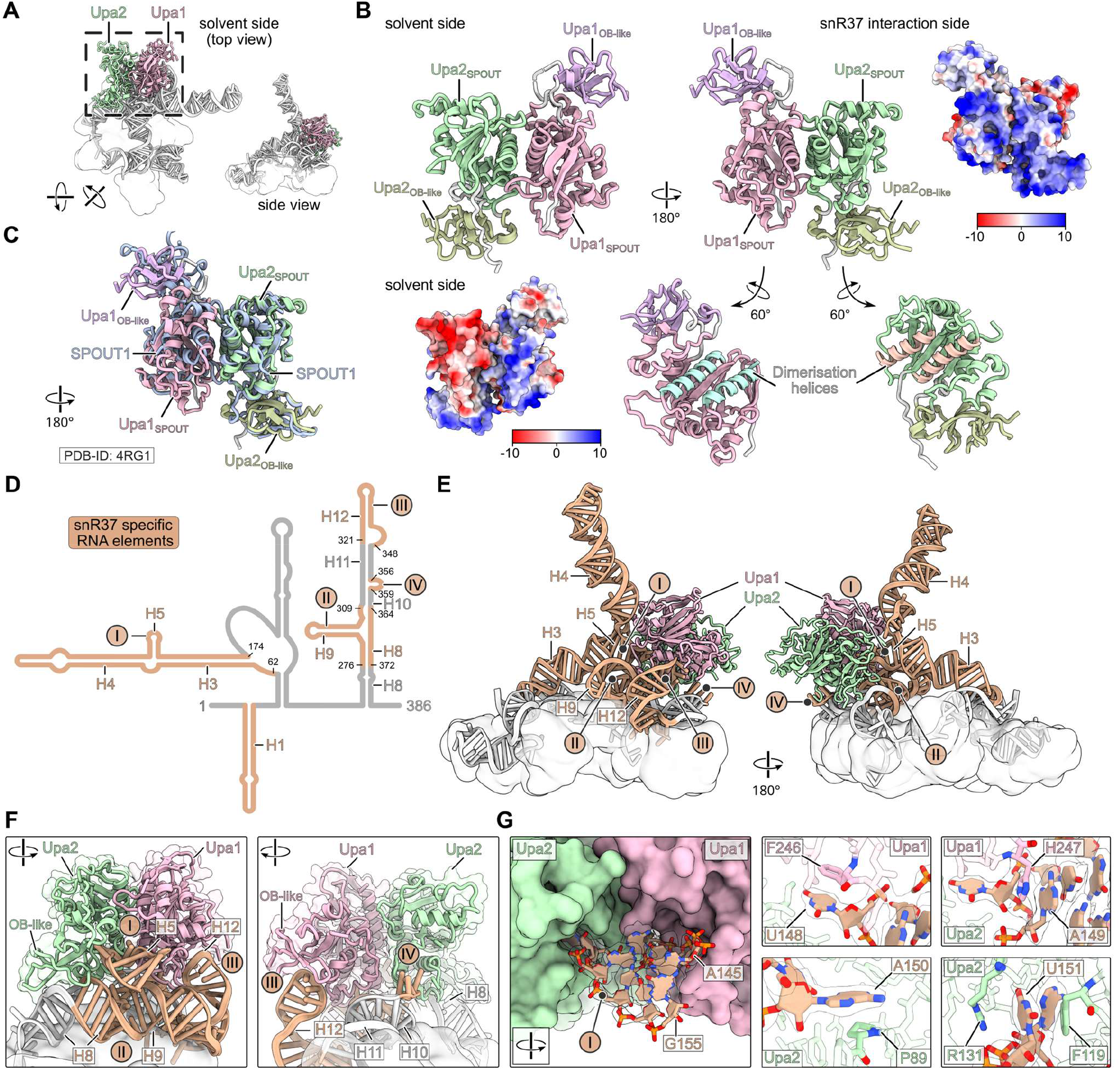
snR37 establishes the interaction platform for Upa1-Upa2, the homologs of human SPOUT1. (**A**) Overviews of the Upa1-Upa2 heterodimer bound to the snR37 snoRNP. (**B**) Solvent side (as shown in A) and snR37-interacting side of the Upa1-Upa2 complex. The SPOUT folds and internal OB-like folds are highlighted. The same views are also shown as surface representation colored according to the electrostatic potential. In the 60° rotated display, the α-helices of Upa1 and Upa2 at the protein-protein interface are highlighted. (**C**) Overlay of the Upa1-Upa2 heterodimer with the homodimer structure of the human homolog SPOUT1 (PDB-ID: 4RG1). (**D**) The secondary structure of snR37 is shown. RNA elements that are specific to the snR37 snoRNA as well as binding sites of Upa1-Upa2, according to the cryo-EM model, are indicated (I-IV) and colored. (**E)** Overview of the snR37 structure focusing on snR37 and its interaction with Upa1-Upa2. The snR37-specific elements are colored and labeled as shown in (D). The H/ACA core proteins are shown as filtered, transparent surfaces for clarity. (**F**) Close-up views of (E) showing magnifications of the snR37-Upa1-Upa2 interaction regions in two orientations. (**G**) The Upa1-Upa2 interaction surface is recognized by snR37 helix 5 (H5, I) through specific stacking interaction between the bases of snR37 residues U148-U151 and the highlighted residues of Upa1 and Upa2. Left panel: Surface view of Upa1-Upa2 together with molecular model and segmented cryo-EM density of snR37 H5. Right panels: Magnified views of the interactions between snR37 nucleotides and Upa1-Upa2 residues.

In our snR37 snoRNP model, Upa1 and Upa2 interact exclusively with the snoRNA and do not contact the H/ACA core proteins (**Figures 4**D, **4**E, **6**D, and **6**E). Specifically, the Upa1-Upa2 heterodimer engages with several unique features of snR37 that are absent from canonical H/ACA snoRNAs (**Figures 6**D and **6**E, I-IV). Here, the OB-like folds of Upa1 and Upa2 interact with the extended upper stem (H12, III) and the lower stem (H8, II) of the 3′ hairpin, respectively (**Figure 6**F). In addition, the highly basic surface of the SPOUT dimer binds the three-way junction formed by H8, H9, and H10 (**Figure 6**B, **6**D, and **6**F). The unique internal 5′ branch harbors two long helices (H3 and H4), which are connected by a short third helix (H5), forming an elbow-like structure (**Figure 6**D and **6**E). Helix H3 extends from the pseudouridylation pocket of the 5′ protomer toward the 3′ hairpin of snR37, positioning helix H5 atop the H8-H9-H10 RNA junction within the 3′ hairpin. This novel conformation enables helix H5 to interact with and recognize the Upa1-Upa2 interaction surface within the 3′ module. Here, the bases of snR37 nucleotides U148-U151 interact specifically with residues of both Upa1 and Upa2 via cation-pi stacking (**Figure 6**D and **6**G).

Although Rbp95 was present in the purified complexes at similar levels as Upa1 and Upa2 (**Figure 4**A) and was even used as bait protein for the Rbp95-Cbf5 split purification (**Figure 4**B), it was not visible in our cryo-EM reconstruction. This is likely due to conformational flexibility or dissociation during sample preparation. Nevertheless, our interaction assays showed that Rbp95 interacts specifically with Upa2 (**Figures 1**F–**1**H).

Taken together, the structural findings are consistent with our biochemical data. The snR37 snoRNP contains the non-canonical proteins Upa1 and Upa2 as a heterodimer that is recruited through unique RNA elements of the snR37 snoRNA. The extensive interaction interface between Upa1-Upa2 and snR37 explains the heterodimer’s binding specificity for the snR37 snoRNA.

### Mutants disentangle catalytic from non-catalytic functions of the snR37 snoRNP

To dissect the different functions of the snR37 snoRNP, we designed a panel of *snr37* mutants informed by known H/ACA snoRNA features as well as insights from our cryo-EM and CRAC data (**Figure 7**A). The engineered snR37 variants included: a single nucleotide deletion predicted to alter the positioning of 25S rRNA residue U2944 in the pseudouridylation pocket and thereby abolish pseudouridylation (ΔC244); a base-pairing mutant in which nine of the sixteen nucleotides of the guide region were replaced by their complementary bases, thus drastically reducing the stability of base pairing with the target rRNA (bp mutant); two truncation variants of the snR37-specific internal 5′ branch (ΔG96-A137 and ΔU78-G162); a truncation variant lacking the 5′ leader sequence (ΔG8-U49); and a variant in which all nucleotides contacting Upa1 and Upa2 according to CRAC and cryo-EM data were deleted or mutated (ΔUpa1/2 sites) (**Figures 7**A and **S10**A). We then assessed steady-state levels and pre-60S association of these snR37 variants (**Figure 7**B), their ability to direct U2944 pseudouridylation (**Figures 7**C and **S10**B), their effects on anisomycin resistance (**Figures 7**D and **S10**C), and their ability to rescue the synthetic growth defects of the *dbp9-5 snr37*Δ and *rpl3-102 snr37*Δ double mutants as an indicator of ribosome biogenesis phenotypes (**Figures 7**E and **S10**D). All variants were expressed, and only the large 5′ stem deletion variant (ΔU78-G162) showed lower levels, suggesting reduced stability (**Figure 7**B, input). Pre-60S association was nearly abolished in the ΔUpa1/2 sites mutant, mirroring the loss of snR37 association in the *upa1*Δ *upa2*Δ background (**Figure 7**B, Rsa3-TAP IP). Surprisingly, all other variants, including the base-pairing mutant lacking the ability to stably interact with the 25S rRNA target site, still co-purified with pre-60S particles at levels comparable to wild-type snR37 (**Figure 7**B). These findings reveal that stable incorporation of snR37 into pre-60S particles requires mainly the interaction with Upa1 and Upa2, but not base pairing of the guide sequence with the 25S rRNA.

**Figure 7.**
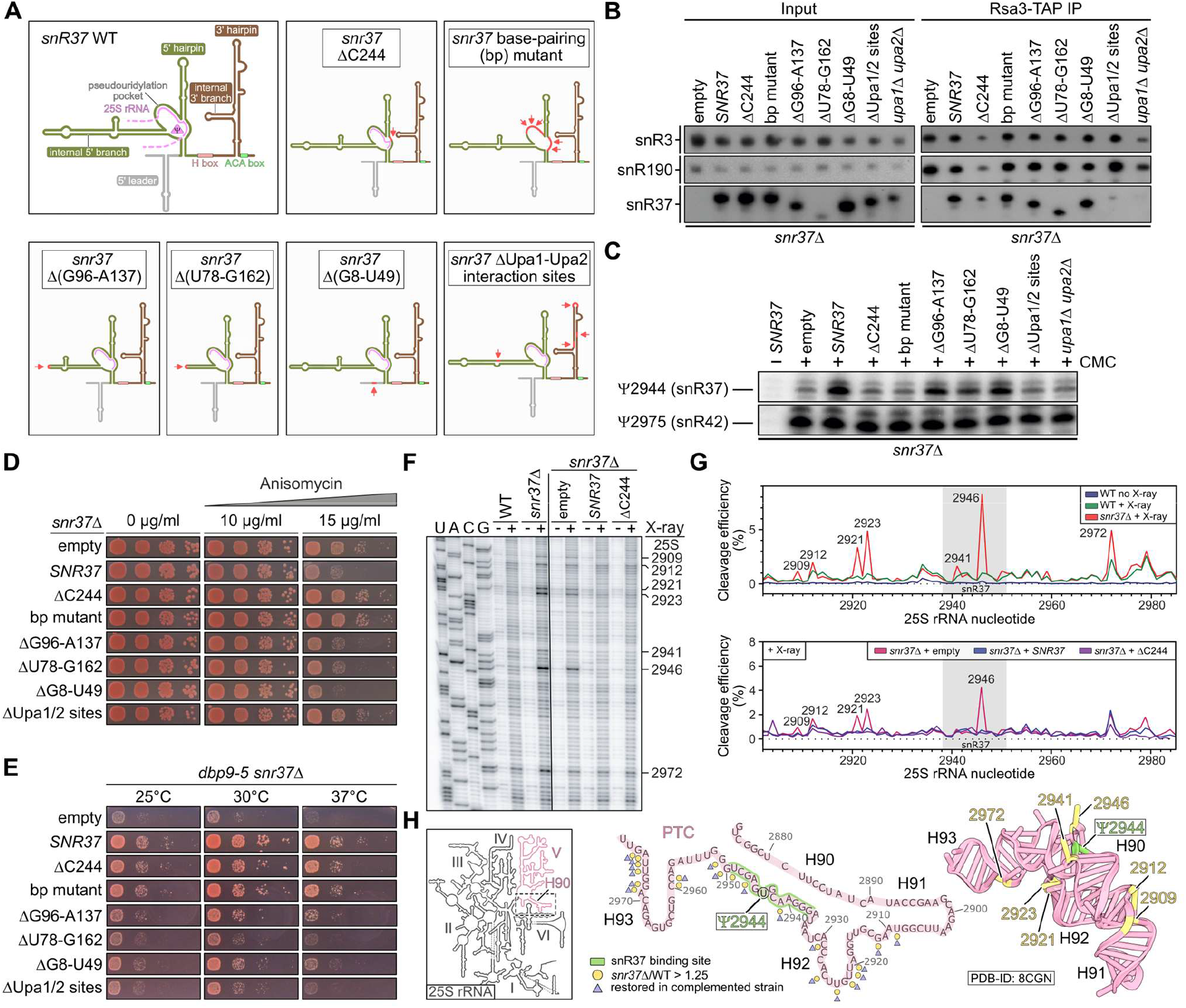
Directed mutants disentangle catalytic from non-catalytic functions of the snR37 snoRNP. (**A**) Schematic overview of wild-type (WT) snR37 and analyzed mutant variants. (**B**) Northern blot analysis of snoRNAs co-purified with Rsa3-TAP pre-60S particles. *snr37*Δ cells transformed with *TRP1* plasmids expressing the indicated variants of snR37 depicted in (A) were compared to wild-type *SNR37, snr37*Δ, and *upa1*Δ *upa2*Δ cells. Probes against snR37, snR190, and snR3 were used. Inputs show snoRNA levels in crude extracts taken prior to purification. (**C**) Pseudouridylation analysis via primer extensions after CMC treatment of RNA from the same strains as in (B). Ψ sites can be identified by primer extension stops in CMC-treated samples. The full primer extension gel is shown in **Figure S10**B. (**D**) *snr37*Δ cells transformed with the same plasmids as in (B) were spotted in serial dilutions on SDC-Trp plates with increasing anisomycin concentrations and incubated at 30 °C for 2 days. (**E**) Growth complementation assay. *dbp9-5 snr37*Δ strains were transformed with the same plasmids as in (B). Serial dilutions were spotted on SDC-Trp plates and incubated for 3 days at the indicated temperatures. **(F)** Sequencing gel showing X-ray-induced cleavage patterns of 25S rRNA after 150 ms exposure in wild-type and *snr37*Δ strains, as well as in *snr37*Δ strains transformed with *TRP1* plasmids harboring no insert (empty vector), *SNR37*, or *snr37*ΔC244. **(G)** X-ray cleavage efficiency plots for a 25S rRNA fragment in wild-type and *snr37*Δ strains (upper panel) and in plasmid-containing *snr37*Δ strains (lower panel). Nucleotides showing increased exposure in *snr37*Δ relative to wild type are indicated. The snR37-binding site is highlighted in gray. **(H)** Structural analysis of 25S rRNA H90-93 based on X-ray footprinting data. In the 2D scheme, the snR37-binding site is shown in green. Yellow circles indicate nucleotides with a *snr37*Δ/WT cleavage efficiency ratio > 1.25, and purple triangles indicate nucleotides whose cleavage patterns were restored in complemented strains. The 3D structure illustrates the spatial relationship between the most perturbed nucleotides in *snr37*Δ strains (yellow) and the Ψ2944 site (green).

We next tested whether the variants retained catalytic function by analyzing pseudouridylation at U2944. As expected, both the ΔC244 and base-pairing mutants failed to modify U2944 (**Figures 7**C and **S10**B). As these variants are still capable of binding to pre-60S (**Figure 7**B), their defect likely reflects an inability to correctly position U2944 for pseudouridylation by Cbf5. Similarly, the ΔUpa1/2 sites mutant failed to mediate pseudouridylation (**Figure 7**C), consistent with its inability to associate with pre-60S particles (**Figure 7**B). In contrast, the 5′ leader truncation and the mutant with the smaller truncation of the 5′ internal branch (ΔG96-A137) largely retained pseudouridylation function. The mutant with the larger 5′ internal branch truncation (ΔU78-G162), which lacks the Upa1-Upa2-interacting helix H5, still supported pseudouridylation, albeit at reduced levels (**Figure 7**C).

Notably, anisomycin resistance was observed in all mutants deficient in U2944 pseudouridylation (ΔC244, bp mutant, and ΔUpa1/2 sites), phenocopying the *snr37*Δ mutant (**Figure 7**D), suggesting that anisomycin resistance is caused by the absence of pseudouridylation. All mutants remained sensitive to the translation inhibitor cycloheximide, confirming that the effects are specific to the PTC-binding drug anisomycin (**Figure S10**C).

Next, we examined whether the snR37 variants could rescue the synthetic growth defects observed in *dbp9-5 snr37*Δ (**Figure 7**E) and *rpl3-102 snr37*Δ (**Figure S10**D) double mutants, where failure to rescue was considered indicative of early pre-60S biogenesis defects. Strikingly, both the ΔC244 and base-pairing mutants, despite being unable to mediate pseudouridylation at U2944 (**Figure 7**C), still complemented the growth defects in both genetic backgrounds (**Figures 7**E and **S10**D). This demonstrates that the snR37 snoRNP has a function in ribosome biogenesis that is independent of its pseudouridylation role and independent of guide sequence-mediated base pairing with the rRNA. By contrast, the ΔUpa1/2 binding sites variant, which fails to stably associate with pre-60S particles (**Figure 7**B), did not rescue the growth phenotypes (**Figures 7**E and **S10**D). Moreover, the three different truncation variants, despite being capable of associating with pre-60S particles (**Figure 7**B) and retaining pseudouridylation function (**Figure 7**C), only partially rescued growth (ΔG96-A137 and ΔG8-U49) or failed to complement (ΔU78-G162) (**Figures 7**E and **S10**D).

Together, these findings reveal that snR37 harbors a non-catalytic function in ribosome biogenesis mediated by its unique RNA elements. To test if snR37 contributes to the correct structuring of the 60S subunit rRNA, we performed X-ray hydroxyl radical probing of total RNA from wild-type and *snr37*Δ cells, focusing on the 25S rRNA region bound by snR37 (**Figures 7**F-**7**H, **S11**A, and **S11**B). In this assay, exposed and flexible RNA regions are preferentially cleaved, producing primer extension stops^59^. Compared with the wild-type situation, the 25S rRNA of *snr37*Δ cells showed prominent cleavage sites in the snR37-binding region in helix H90 and in the adjacent helices H92 and H93, indicating a distortion of the PTC (**Figures 7**F-**7**H). In contrast, an 18S rRNA region tested as control showed no structural differences (**Figure S11**C). Given that snR37 acts on early pre-60S precursors and that the altered structure was detected in total RNA, the observation of structural distortion in the PTC is remarkable. This suggests that loss of snR37 binding during early pre-60S assembly has lasting structural consequences that persist in mature ribosomes.

We next asked whether the difference arises from the presence of uridine (U2944) in *snr37*Δ cells versus pseudouridine (Ψ2944) in wild-type cells. Footprinting results of *snr37*Δ cells transformed with either empty plasmid, plasmid expressing wild-type *SNR37*, or the pseudouridylation deficient mutant *snr37*ΔC244 showed that both wild-type and ΔC244 snoRNA restored the structure (**Figures 7**F-**7**H, and **S11**B). Thus, the observed PTC distortion is independent of pseudouridylation, revealing a role of snR37 in rRNA folding.

In summary, we demonstrate that snR37 is an H/ACA snoRNP with dual functionality: while its canonical guide sequence directs site-specific pseudouridylation that influences anisomycin resistance, its additional structural elements perform an independent, non-catalytic role in ribosome biogenesis that supports the assembly process and is essential for establishing the correct local architecture of the PTC.

## Discussion

In this study, we present the structure of a bifunctional eukaryotic box H/ACA snoRNP that functions in both rRNA modification and rRNA scaffolding. Our data reveal that snR37 contains a canonical 5′ hairpin bound by the four H/ACA core proteins Cbf5, Nhp2, Nop10, and Gar1, forming a functional pseudouridylation module that modifies 25S rRNA residue U2944 in the A site of the PTC. In contrast, the 3′ hairpin of snR37 adopts a non-canonical configuration lacking a pseudouridylation pocket that is bound by Cbf5, Nhp2, and Nop10 but lacks Gar1. Instead, the typical position of Gar1 is occupied by the snR37-specific internal 3′ branch. The Upa1-Upa2 heterodimer interacts with unique RNA features of snR37, explaining why these specialized proteins are recruited to snR37 but not other H/ACA snoRNAs. Biochemical evidence confirms that Rbp95 is also part of the snR37 snoRNP, although it could not be visualized in the cryo-EM structure, likely due to conformational flexibility or loss during cryo-EM sample preparation.

Interestingly, both Upa1 and Upa2 harbor a SPOUT domain, a hallmark of a diverse family of RNA and protein methyltransferases that typically function as homodimers^44^. Upa1 and Upa2 however form a heterodimer in which the individual subunits fulfill distinct roles: Upa2 mediates the interaction with Rbp95, whereas Upa1 connects the snoRNP to the Npa1 complex.

Structural superimposition of yeast Upa1-Upa2 with human SPOUT1, a catalytically active homodimer,^45, 57^ indicates that the yeast heterodimer is catalytically inactive, as neither subunit appears able to bind the methyl donor SAM when associated with the snR37 snoRNP. Accordingly, whereas Drosophila homolog Ptch and human SPOUT1 methylate residues in the PTC (U3485 in flies, U4530 in humans), the corresponding site in yeast (U2953) remains unmodified.^58^ The residue pseudouridylated by yeast snR37 (U2944) corresponds to U4521 in humans, predicted to be modified by SNORA10 (**Figure S9**D).^60, 61^ The close spatial proximity of U4521 to the SPOUT1-methylated U4530 suggests that, similar to yeast Upa1-Upa2, SPOUT1 may, beyond its catalytic role, also contribute structurally to 60S subunit maturation, potentially in concert with SNORA10. This structural function could be relevant for disease, as patient-derived SPOUT1 variants are associated with severe neurodevelopmental disorders.^62^

A previous study reported that deletion of *SNR37* increases the accuracy of stop codon recognition, suggesting that snR37 fine-tunes translation fidelity.^49^ We observed two distinct types of lasting alterations in the PTC region. The first arises when snR37’s pseudouridylation function is lost, leading to resistance to anisomycin, an inhibitor of aminoacyl-tRNA binding that targets the A site of the PTC.^50^ This resistance suggests subtle remodeling of the A-site environment in the absence of U2944 modification. The second structural alteration, revealed by hydroxyl radical footprinting, is caused by loss of the modification-independent function of snR37 and manifests as distortions in 25S rRNA helices H90, H92, and H93. This indicates that the scaffolding function of snR37 is required to establish the correct architecture of this region.

We propose that, beyond these local effects at its base-pairing site, snR37 also contributes to higher-order organization of pre-60S particles. The synthetic growth and ribosome biogenesis defects of the *snr37*Δ mutant upon combination with mutations in genes of the *NPA1* genetic network suggests a partially redundant function of snR37 and the Npa1 complex. Truncation analysis further revealed that the 5′ internal branch and the 5′ leader of snR37 are critical for this function. The precise nature of this non-catalytic role in ribosome-biogenesis remains elusive, but it is likely linked to the extensive interaction network in which snR37 engages: it base-pairs with helix H90 in domain V, contacts helix H95 in domain VI via Rbp95, and interacts with the Npa1 complex, which itself binds domains I, II, and VI^31, 63^. It is therefore highly plausible that the snR37 snoRNP, in cooperation with the Npa1 complex, functions in rRNA folding by tethering and sterically coordinating distinct rRNA domains during early stages of 60S subunit assembly. In this early phase of the 60S assembly pathway, clamping the most distant rRNA regions (domain I/II and domains V/VI) together would be particularly advantageous to prime the overall compaction process in a sterically correct manner. This model is consistent with the observation that, apart from portions of domain V, these distal domains are already structured within the earliest resolved nucleolar pre-60S intermediates.^64, 65^ Interestingly, another atypical snoRNA, snR190, also interacts with the Npa1 complex,^23, 28^ suggesting that it is also part of this intricate chaperoning network.

In addition to snR190, two other yeast snoRNPs, U3 and snR30, do not execute any rRNA modification, but instead serve essential structural or chaperoning roles in ribosome assembly. It has been believed that such “assembly” snoRNPs represent rare exceptions.^66, 67^ Importantly, our findings reveal that chaperone functions in ribosome assembly are not reserved to snoRNPs without catalytic function, but can also be performed by snoRNPs active in rRNA modification. Indeed, two recent studies in fission yeast and humans describe bifunctional snoRNAs. The fission yeast box C/D snoRNA snR107 directs 2′-*O*-ribose methylation of 25S rRNA, but also associates with gametogenic RNA-binding proteins to repress meiotic gene expression.^68^ Similarly, human SNORA13 not only catalyzes pseudouridylation but additionally sequesters ribosomal protein uL14 (RPL23), thereby inhibiting ribosome biogenesis and promoting senescence.^69^ The precise mechanisms of these additional interactions, including whether additional protein components are part of these unconventional snoRNPs, remain to be elucidated.

In *S. cerevisiae*, all 2′-*O*-ribose methylation and pseudouridylation sites have been mapped and assigned to specific snoRNAs. Eukaryotic H/ACA snoRNAs universally adopt a bipartite structure with two hairpins, hence theoretically having the potential to guide two modifications. However, according to the UMass-Amherst Yeast snoRNA Database,^47^ half of the 28 yeast H/ACA snoRNAs mediate only a single modification, raising the possibility that their modification-inactive hairpins serve alternative roles. Our results on the dual function of snR37 suggest that the current dichotomy of snoRNPs falling into two classes with either catalytic or structural/chaperoning functions may be too simplistic. Instead, our finding argues for a continuum of snoRNP function, where dual function, combining rRNA modification and structural/chaperoning roles, may be a more prevalent principle rather than an exception.

## Methods

### Yeast strains and plasmid construction

All *S. cerevisiae* strains used in this study were generated by chromosomal deletion and chromosomal tagging using established methods.^70, 71^ All strains are listed in **Table S1**. All plasmids used in this study for *E. coli* and *S. cerevisiae* were constructed using standard DNA cloning techniques and they are listed in **Table S2**. All fragments that were amplified by PCR were validated by sequencing.

### Fluorescence Microscopy

Yeast cells expressing C-terminally GFP-tagged Upa1 or Upa2 as well as the nucleolar marker protein Nop58-RedStar from the genomic locus were imaged in logarithmic growth phase using a Leica DM6 B microscope with GFP or RHOD ET filters, a × 100/1.4 Plan APO objective, a DFC 9000 GT camera and the LasX software.

### Yeast two-hybrid (Y2H)

For analysis of protein-protein interactions, Y2H assays using the reporter strain PJ69-4A were performed. The reporter strain was transformed with two different plasmids, one carrying the bait protein, fused to the DNA-binding domain of the Gal4 transcription factor (G4BD with a c-Myc tag, BD, *TRP1* marker), and the other one carrying the prey protein, fused to the activation domain of Gal4 transcription factor (G4AD with an HA tag, AD, *LEU2* marker). The G4BD or G4AD were either fused C-terminally (ADC or BDC) or N-terminally (ADN or BDN) to bait or prey proteins. Transformed yeast cells carrying both plasmids were spotted in a serial dilution on plates lacking leucine and tryptophan (SDC-Leu-Trp, control plates), lacking leucine, tryptophan, and histidine (SDC-Leu-Trp-His, *HIS3* reporter gene, for detection of weak protein-protein interactions), and plates lacking leucine, tryptophan, and adenine (SDC-Leu-Trp-Ade, *ADE2* reporter gene, for detection of strong protein-protein interactions) and incubated for 3-4 days at 30 °C.

### Drug sensitivity assays

Strains YPH499, YHH1, YHH2, and YHH3 (**Table S1**) were cultivated at 30 °C in YPD to exponential phase (OD_600_ = 0.5). Cells were harvested by centrifugation, washed twice with sterile water, and resuspended to an OD_600_ of 0.1. Ten-fold serial dilutions were prepared in sterile water and 3 µl drops were spotted onto control YPD plates (2% agar) and plates containing anisomycin (from 2 to 10 µg/ml; MilliporeSigma) or cycloheximide (0.5 µg/ml; MilliporeSigma). Plates were incubated at 30 °C for 2 to 5 days.

To test drug sensitivity of different *snr37* mutants, *snr37*Δ yeast strains were transformed with *TRP1* plasmids expressing wild-type snR37 (WT), different snR37 variants (ΔC244, bp mutant, ΔG96-A137, ΔU78-G162, ΔG8-U49, ΔUpa1/2 sites) or were transformed with a *TRP1* empty plasmid to reflect a *SNR37* knockout (*snr37*Δ). Transformed cells were spotted in a serial dilution on SDC-Trp plates with increasing anisomycin concentration (0, 10 or 20 µg/ml; Sigma-Aldrich #A5862) and incubated for 3-4 days at 30 °C or with increasing cycloheximide concentrations (0, 0.03 or 0.3 µg/ml; Sigma-Aldrich #C7698) and incubated for 2 days at 30 °C.

### Polysome profiling

Yeast cells carrying a knockout of *UPA1* and *UPA2* were used. Experiments were performed as described previously.^23^

### TurboID-based proximity labeling

Plasmids expressing N-or C-terminally TurboID-tagged Upa1 and Upa2 under the control of the cooper-inducible *CUP1* promoter were transformed into the wild-type strain YDK11-5A. The experiment was performed as described previously.^36, 72^

### Heterologous expression and co-purification assays of Upa1, Upa2, and Rbp95

Genes encoding (His)_6_-Upa1 (MCS1), Flag-Upa2 (MCS2), (His)_6_-Rbp95 (MCS1), and Flag-Upa1 (MCS2) on their own, as well as combinations of (His)_6_-Upa1 with Flag-Upa2, (His)_6_-Rbp95 with Flag-Upa1, and (His)_6_-Rbp95 with Flag-Upa2, were cloned into pETDuet-1 (Novagen). Plasmids were transformed into the *E. coli* BL21 (DE3) Rosetta strain (Novagen). Cells were grown in 2 l 2x TY media at 37 °C to an OD_600_ of 0.3 and then protein expression was induced using 0.3 mM isopropyl-β-D-thiogalactoside (IPTG, Thermo Scientific). Cells were then further incubated at 16 °C for 20 h and were then harvested. Pellets were resuspended in lysis buffer (50 mM Na_2_HPO_4_/NaH_2_PO_4_ buffer pH 7.5 with 150 mM NaCl) containing 1x Protease Inhibitor Mix HP (Serva), 0.5 mM phenylmethanesulfonyl fluoride (PMSF, Sigma-Aldrich), 1 mM dithiothreitol (DTT, Roth), 1 mg/ml lysozyme (Roth) and incubated for 40 min on ice. Cells were lysed by sonication and after centrifugation cell lysates were treated with RNase A (NEB, 20 mg/ml) for 15 minutes at 25 °C and after that incubated with 300 µl of Anti-FLAG^®^ M2 Affinity Gel (Sigma-Aldrich) under rotation for 1 h at 4 °C. Beads were washed three times with lysis buffer containing 1 mM DTT, transferred to Mobicol columns (MoBiTec) and washed with additional 5 ml of buffer. Washed beads were incubated with 300 µl of lysis buffer containing 1 mM DTT and 100 µg/ml 1xFLAG peptide (Sigma-Aldrich) under rotation for 1 h at 4 ° C for elution of bound proteins. Imidazole was added to eluates to a final concentration of 60 mM. Eluates were then incubated with 300 µl of Ni-NTA agarose (Qiagen) under rotation for 1 h at 4 °C. Beads were washed three times with lysis buffer containing 1 mM DTT and 60 mM imidazole and after that transferred to Mobicol columns and washed with additional 5 ml of buffer. For elution of bound proteins, washed beads were incubated with lysis buffer containing 1 mM DTT and 300 mM imidazole under rotation for 1 h at 4 °C. Samples were analyzed via NuPAGE^™^ 12% Bis-Tris gels (Invitrogen) followed by western blotting or staining with NOVEX Colloidal Blue Staining kit (Invitrogen).

### CRAC experiment and analysis

The CRAC experiment was performed using yeast cells expressing Upa1 and Upa2 C-terminally fused to a HTP tag (consisting of a (His)_6_ tag, TEV protease cleavage site, and two Z-domains of protein A) as well as a wild-type untagged strain as control. Experiments and data analysis were performed as described previously.^36^

### Immmunoprecipitation experiments with Upa1-Myc and Upa2-HA and western blotting

Strain YHH4 (**Table S1**) was cultivated at 30 °C in YPD to exponential phase (OD_600_ = 0.5). Cells were harvested by centrifugation and washed twice with sterile water. The equivalent of 10 OD_600_ units of cells were used for each immunoprecipitation experiment. Whole cell extracts were prepared in TMN100 buffer (25 mM Tris-HCl pH 7.9, 10 mM MgCl_2_, 100 mM NaCl, 0.1% NP-40, 1 mM DTT, supplemented with cOmplete™ protease inhibitor cocktail (Roche)) by mechanical disruption with glass beads, as described.^73^ Cell lysates (500 µl) were incubated with anti-HA or anti-myc magnetic beads (MedChemExpress) for 2 h at 4 °C. Then, beads were recovered with a magnet, washed twice with TMN100, resuspended in 50 µl 2x Laemmli sample buffer, and incubated at 95 °C for 5 min. Eluted proteins were fractionated by SDS-PAGE and analyzed by western blotting. Bands were visualized with a Fusion FX7 imaging system.

### Tandem affinity purification (TAP)

For TAP purification, yeast cells expressing C-terminally TAP (Tandem Affinity Purification tag, consisting of C-terminal calmodulin-binding peptide, TEV cleavage site, and two Z-domains of protein A)-tagged fusion proteins were grown in 4 L YPD to an OD_600_ of 2. TAP purifications were performed in a buffer containing 50 mM Tris-HCl pH 7.5, 100 mM NaCl, 1.5 mM MgCl_2,_ 0.1% NP-40, 1 mM DTT and freshly added 1x Protease Inhibitor Mix FY (Serva). Cells were lysed by mechanical disruption using glass beads. Cell lysates were cleared by centrifugation and incubated with 450 µl of IgG Sepharose^™^ 6 Fast Flow (GE Healthcare) under rotation for 1 h at 4 °C. Beads were then transferred into Mobicol columns (MoBiTec) and washed with 10 ml of buffer. For elution of bound proteins, the beads were incubated with 300 µl of buffer containing TEV protease in the presence of 100 U RiboLock RNase inhibitor (Thermo Scientific) under rotation for 70 min at room temperature. CaCl_2_ was added to the TEV eluate to a final concentration of 2 mM and then the eluate was incubated with 300 µl of Calmodulin Sepharose^™^ 4B (GE Healthcare) under rotation for 1 h at 4 °C. Beads were then washed with 5 ml of buffer containing 2 mM CaCl_2._ After washing, elution was performed with 600 µl buffer containing 5 mM EGTA for 20 min at RT. Eluates were then either used for RNA extraction, or proteins were precipitated with trichloroacetic acid (10% TCA). Final samples were separated on NuPAGE^™^ 4-12% Bis-Tris gels (Invitrogen) followed by staining with NOVEX^®^ Colloidal Blue Staining Kit (Invitrogen) or western blotting.

### Immunoprecipitation experiments of Upa1-TAP, Upa2-TAP, or Prp22-TAP and RNA extraction

Cell pellets from strains expressing Upa1-TAP, Upa2-TAP or Prp22-TAP were resuspended with ice-cold A200-KCl buffer (20 mM Tris-HCl pH 8.0, 5 mM magnesium acetate, 200 mM KCl, 0.2% Triton-X100) supplemented with 1 mM DTT, 1x cOmplete^™^ EDTA-free protease inhibitor cocktail (Roche), 0.1 U/µl RNasin (Promega) and cells were broken by vigorous shaking using zirconia beads. Extracts were clarified at 16,000 × g and 4 °C for 10 min and equal amounts of soluble extracts in a final volume of 1 ml were incubated for 2 h at 4 °C with IgG-Sepharose beads on a rocking table. Beads were washed seven times with 1 ml of ice-cold A200-KCl buffer supplemented with 1 mM DTT. RNA was extracted from bead pellets as follows: 160 µl of 4 M guanidium isothiocyanate solution, 4 µl of glycogen (Roche), 80 µl of (10 mM Tris-HCl pH 8.0, 1 mM EDTA, pH 8.0, 100 mM sodium acetate), 120 µl of phenol and 120 µl of chloroform were added. Tubes were shaken vigorously, incubated 5 min at 65 °C, and centrifuged 5 min at 16,000 × g (4 °C). Aqueous phases (240 µl) were mixed vigorously with 120 µl of phenol and 120 µl of chloroform, centrifuged 5 min at 16,000 × g (4 °C) and the resulting aqueous phases were ethanol precipitated.

### Primer extension experiments

Primer extension experiments to detect rRNA processing intermediates were performed as follows. RNAs extracted from total input extracts or from samples obtained following precipitation of TAP-tagged proteins using IgG-Sepharose were denatured at 80 °C during 4 min and annealed to ^32^P-labelled oligonucleotides (5′-CTCAATACGCATCAACCCATGTC-3′ to detect 35S pre-rRNA and 5′-GGCCAGCAATTTCAAGTTA-3′ to detect 27SA_2_ and 27SB pre-rRNAs) at 50 °C during 90 min in 0.3 M NaCl, 10 mM Tris-HCl pH 7.5, 2 mM EDTA pH 8.0. Primer extensions were then carried out with AMV reverse transcriptase (Promega) in 1 mM dATP/dCTP/dGTP/dTTP, 10 mM DTT, 10 mM Tris-HCl pH 8.4, 6 mM MgCl_2_ for 1 h at 42 °C. The reactions were stopped by addition of 10 mM final EDTA, pH 8.0. RNAs were hydrolyzed in 0.1 M NaOH at 55 °C for 1 h. NaOH was neutralized with HCl. The cDNAs were ethanol precipitated, resuspended in H_2_O and subjected to denaturing electrophoresis in 6% acrylamide/urea sequencing gels in TBE as running buffer.

Primer extension experiments to detect 2′-*O*-ribose methylation were performed using yeast cells carrying a knockout of *UPA1* and *UPA2, SNR37, SNR69, SNR71*, or *SNR73*. RNA isolated from these cells were mixed with 0.2 pmol of radiolabeled primer, which binds downstream of modification sites. After denaturation (95 °C, 1 min), primer extension was performed with 10 units of AMV reverse transcriptase (Promega) according to the manufacturer’s protocol either with 1 mM dNTPs (high concentration) or 0.1 mM dNTPs (low concentration). The cDNAs were then separated by denaturing electrophoresis in 6% acrylamide/urea sequencing gels in TBE as running buffer and analyzed using a phosphoscreen.

Primer extension experiments to detect pseudouridines (Ψ) were performed using yeast cells carrying a knockout of *UPA1* and *UPA2* or *SNR37*. RNA extracted from these cells was treated with CMC, which binds to uridines and pseudouridines, followed by alkaline treatment, which removes CMC from uridines but not pseudouridines. Primer extension experiments were performed with regular dNTP concentrations and with 10 units of AMV reverse transcriptase (Promega) according to the manufacturer’s protocol. The cDNAs were then separated by denaturing electrophoresis in 6% acrylamide/urea sequencing gels in TBE as running buffer and analyzed using a phosphoscreen.

### Northern Blotting

For total RNA isolation, yeast cells were grown in 20 ml YPD at 25 °C or 30 °C to an OD_600_ of ∼0.5. Cell pellets were resuspended in 200 µl lysis buffer (10 mM Tris-HCl pH 7.5, 10 mM EDTA, 0.5% SDS) and mechanically disrupted using 200 µl glass beads. Cell lysates were centrifuged and supernatants were used for RNA isolation. For RNA isolation from TAP purification samples, 20 µl of cell lysates were used and adjusted to 600 µl volume using TAP elution buffer or 600 µl of final TAP eluates were used.

RNA was extracted by using two phenol-chloroform-isoamyl alcohol extractions (25:24:1) and one chloroform-isoamyl alcohol (24:1) extraction. RNA precipitation was performed with addition of 1/10 volume of 3 M sodium acetate pH 5.2, 2.5 volumes of 100% ethanol, and 1 µl of GlycoBlue^™^ (Invitrogen). RNA pellets were dissolved in nuclease-free water. For analysis of pre-rRNAs, 1-3 µg of extracted RNAs were separated on 1.6% agarose gels containing MOPS buffer (20 mM 3-(N-morpholino)-propanesulfonic acid (MOPS) pH 7.0, 5 mM sodium acetate, 1 mM EDTA, 0.75% formaldehyde) and ethidium bromide and electrophoresis was carried out at 60 V for 6-8 h in MOPS buffer. RNAs were transferred onto Hybond N^+^ nylon membranes (Amersham) by capillary transfer overnight and UV crosslinked to the membrane. For detection of snoRNAs, RNA samples were separated on NuPAGE^™^ 6% TBE-Urea gels (Invitrogen). Blotting was performed onto Hybond N^+^ nylon membranes using 0.5x TBE buffer at 300 mA and transferred RNA was UV crosslinked to the membrane. For detection of different snoRNAs and pre-rRNA processing intermediates, the following 5′-^32^P-radiolabeled oligonucleotides were used: E/C_2_ (27S A+B): 5′-GGC CAG CAA TTT CAA GTT A-3′; D/A_2_ (20S): 5′-GAC TCT CCA TCT CTT GTC TTC TTG-3′; A_2_A_3_ (35S, 32S/33S, 27SA_2_, 23S): 5′-TGT TAC CTC TGG GCC C-3′; 18S: 5′-GCA TGG CTT AAT CTT TGA GAC-3′; 25S: 5′-CTC CGC TTA TTG ATA TGC-3′; snR37 5′-CAG GGA ATT TCC TCA GAA TAA TAT TCG ATC-3′; snR190 5′-CCTTGTCGTCATGGTCGAATCG -3′; snR69 5′-CCT TTC ATT AAA AAC GAA TCG AAG AGC TGG G-3′; snR71 5′-CAT ATC AAA AGA TCT GAG TGA GCT GAG AAG G -3′; snR73 5′-GCT CAG TAC CAC GCC CTG TCA CAG GCG -3′; snR30 5′-GGC AAC AGC CCC CGA ACC CCA TAT ACA CAT CG -3′, snR10 5′-CCT TGT CGT CAT GGT CGA ATC G -3′

Hybridization was performed with radiolabeled probes under rotation overnight in buffer containing 0.5 M Na_2_HPO_4_ pH 7.2, 7% SDS, and 1 mM EDTA either at 37 °C for the E/C2 probe or 42 °C for all other probes. After washing the membranes four times with buffer containing 40 mM Na_2_HPO_4_ pH 7.2, 1% SDS, radiolabeled signals were detected by using light sensitive X-ray films. Prior to hybridization with another radiolabeled probe membranes were regenerated by washing with 1% SDS for four times at 65 °C.

### Western Blotting

Proteins separated by SDS-PAGE (see above) were transferred to a PVDF membrane in CAPS buffer (10 mM CAPS, 10% methanol) for 2 h with 220 mA. Western blot analysis was performed using the following antibodies: α-His (1:10,000; Sigma-Aldrich, cat. no. A7058); α-Flag (1:15,000; Sigma-Aldrich, cat. no. A8592); α-Rbp95 (1:5,000) generated against full-length recombinant Rbp95 in rabbits ^36^; α-Upa1 generated in this study against full-length recombinant Upa1 in rabbit (1:1,000) (Eurogentec); α-CBP (1:5,000; Merck-Millipore, cat.no. 07-482); α-Dbp6 (1:10,000), α-Npa1 (1:5,000) and α-Rsa3 (1:10,000) ^31^; α-Nhp2 (1:5,000)^74^; α-Prp43 (1:4,000) ^75^; α-Nop1 (1:30,000); α-Noc2 (1:5,000), provided by Herbert Tschochner; α-Rpl3 (1:5,000), provided by Jonathan Warner; α-Rpl16 (1:40,000), provided by Sabine Rospert; secondary α-rabbit horseradish peroxidase-conjugated antibody (1:15,000; Sigma-Aldrich, cat. no. A0545); secondary α-mouse horseradish peroxidase-conjugated antibody (1:10,000; Sigma-Aldrich, cat. no. NA931). Protein signals were visualized using the Clarity^™^ Western ECL Substrate Kit (Bio-Rad) and captured by ChemiDoc^™^ Touch Imaging System (Bio-Rad).

### Genetic interaction assays

*DBP6, DBP9*, and *RPL3* shuffle strains carrying the respective wild-type gene on a *URA3* plasmid were kindly provided by Jesús de la Cruz. To analyze genetic interactions, *UPA1, UPA2*, and *SNR37* were knocked out in these shuffle strains. Strains were transformed with plasmids carrying different alleles of *DBP6, DBP9*, and *RPL3* (*LEU2* marker). The ability of transformants to grow after loss of the respective wild-type gene (on *URA3* plasmid) on plates containing 5-FOA (Thermo Scientific) was evaluated. Transformants that were able to grow on plates containing 5-FOA were further analyzed for growth phenotypes on SDC-Leu plates, which were incubated at for 3 days (25 °C, 30 °C, and 37 °C) or 6 days (16 °C). To analyze genetic interaction between *UPA1, UPA2, SNR37, RBP95* and *RSA3* the respective genes were knocked out in different strains and tested for growth phenotypes on SDC plates, which were incubated for 3 days (25 °C, 30 °C, and 37 °C) or 6 days (16 °C).

### Upa1-FTpA affinity purification and sucrose gradient fractionation

Yeast cells expressing C-terminally FTpA (consisting of a Flag tag, TEV cleavage site, and two Z-domains of protein A)-tagged Upa1 were grown at 30 °C and harvested at an OD_600_ of 2.3. Lysis was carried out using a cryogenic mill (Retsch MM400). The cryo-milled cell powder was resuspended in purification buffer (50 mM Tris-HCl pH 7.5, 100 mM NaCl, 5 mM MgCl_2_, 5% glycerol, 0.1% IGEPAL CA-630, and 1 mM DTT) supplemented with SIGMAFAST protease inhibitor cocktail (Sigma-Aldrich) and RiboLock RNase inhibitor. The lysate was pre-cleared by centrifugation at 4 °C for 10 min at 5,000 rpm, followed by an additional 20 min centrifugation at 17,000 rpm. For the first affinity purification step, the clarified supernatant was incubated with pre-equilibrated IgG Sepharose 6 Fast Flow resin (GE Healthcare) for 2.5 h at 4 °C. The resin was subsequently washed with purification buffer. The resin was collected, and bound proteins were eluted by cleavage with TEV protease at 16 °C for 75 min. For the second affinity purification step, the TEV eluate was incubated with pre-equilibrated anti-FLAG^®^ M2 Affinity Gel (Sigma-Aldrich) for 1.5 h at 4 °C. The resin was then washed with purification buffer, and bound proteins were eluted using 3x FLAG peptide (final concentration of 300 µg/ml) for 45 min at 4 °C.

The Flag eluate of affinity purified Upa1-FTpA was loaded on a 10-40% (w/v) linear sucrose gradient (prepared in buffer containing 100 mM NaCl, 5 mM MgCl2, 50 mM Tris-HCl pH 7.5) and centrifuged at 27,000 rpm for 16 h at 4 °C. Following centrifugation, 13 fractions were collected. One-quarter of each fraction was used for RNA extraction using phenol–chloroform, while the remaining three-quarters were precipitated with 10% TCA. The TCA-precipitated proteins were resuspended in SDS sample buffer and analysed on 4–12% polyacrylamide gels (NuPAGE, Invitrogen), followed by colloidal Coomassie staining. For detection of snR37, extracted RNA samples were resolved on an 8% polyacrylamide/8 M urea gel. RNA was visualized by incubating the gel for 30 min with SYBR Green II RNA gel stain (Sigma–Aldrich) at a 1:5,000 dilution in 1x TBE.

### Mass spectrometry analysis of gel bands

The gel bands were excised and destained before being washed with 50 mM NH_4_HCO_3_. The proteins were reduced by incubating the bands with 45 mM dithioerythritol (DTE) at 55 °C for 30 min and then alkylated with 100 mM iodoacetamide in the dark at room temperature. Following alkylation, the gel pieces were washed again with 50 mM NH_4_HCO_3_. In-gel digestion was performed overnight at 37 °C using 70 ng of sequencing-grade modified trypsin (Promega, Germany). The peptide-containing supernatants were collected and the peptides extracted using 70% acetonitrile (ACN). The fractions were pooled and dried using a vacuum centrifuge (Bachofer, Germany). Protein identification was performed with an Ultimate 3000 nano-liquid chromatography system connected to a Q Exactive HF-X mass spectrometer (both Thermo Scientific). For this, peptides were dissolved in 15 µl of 0.1% formic acid and injected onto an Acclaim PepMap 100 trap column (nanoViper C18, 2 cm length, 100 μM ID, Thermo Scientific). Separation was performed with an analytical column (Aurora XT C18, 25 cm length, 75 μm ID, IonOpticks, Collingwood, Australia) at a flow rate of 250 nl/min. Solvent A was 0.1% formic acid and as solvent B 0.1% formic acid in acetonitrile was used. For separation a 30 min gradient from 3% to 25% solvent B followed by a 5 min gradient from 25% to 40% B was used. Peptides were identified with an online coupled Q Exactive HF-X mass spectrometer (Thermo Scientific). Data-dependent mass spectrometry was performed using cycles of one full MS scan (350-1600 m/z) at 60k resolution and up to 12 MS/MS scans at 15k resolution. The spectra were searched using MASCOT V3.1 (Matrix Science Ltd, London, UK) against the yeast subset of the UniProt database.

### Cryo-EM sample preparation

The Upa1-TAP Cbf5-Flag (Upa1-Cbf5) and Rbp95-TAP Cbf5-Flag (Rbp95-Cbf5) strains were grown in 12 l of YPD medium until an OD_600_ of around 2.5. Then, cells were harvested by centrifugation, frozen in liquid nitrogen, and lysed using a SPEX SamplePrep 6970EFM Freezer/Mill. The cell powder was subsequently stored at -80 °C and was then, prior to affinity purification, resuspended in buffer (60 mM Tris-HCl pH 7.5, 50 mM NaCl, 40 mM KCl, 5 mM MgCl_2_, 1 mM DTT) supplemented with 5% glycerol and 0.1% NP-40, and cOmplete EDTA-free protease inhibitor (Roche). The lysate was pre-cleared by centrifugation at 4,000 rpm for 15 min (Eppendorf centrifuge 5810 R), and the supernatant was centrifuged again at 17,500 rpm for 25 min at 4 °C (Sorvall LYNX 6000 superspeed centrifuge). IgG Sepharose 6 Fast Flow resin (Cytiva) was added to the supernatant, which was then incubated for 90 min on a turning wheel at 4 °C. After centrifugation at 1,800 rpm (Eppendorf centrifuge 5810 R), the supernatant was removed, and the resin was washed once in batch, followed by a wash step using a 20 ml Econo-Pac chromatography column (Bio-Rad). The resin was collected and resuspended in buffer supplemented with 5% glycerol, 0.1% NP-40, and homemade TEV protease. Cleavage was carried out for 90 min at 16 °C under rotation. The supernatant was collected and incubated with anti-FLAG^®^ M2 Affinity Gel (Sigma-Aldrich) for 90 min at 4 °C on a rotating wheel. The beads were collected and washed in batch using 4 ml of buffer supplemented with 2% glycerol and 0.01% NP-40, followed by a second wash with 4 ml of elution buffer (60 mM Tris pH 7.5, 50 mM NaCl, 40 mM KCl, 5 mM MgCl_2_, 1 mM DTT, 2% glycerol, and 0.05% octaethylene glycol monododecyl ether (Nikkol)). The beads were transferred to a 1 ml gravity-flow column (Mobicol) and finally washed with 5 ml of the same buffer. The samples were incubated with elution buffer supplemented with 3xFLAG peptide (Sigma-Aldrich, 250 μg/ml final concentration) for 60 min at 4 °C with shaking, after which the samples were eluted by centrifugation at 2,000 rpm for 20 sec in a tabletop centrifuge at 4 °C.

### Cryo-EM data acquisition and collection

Grid freezing was performed using sample concentrations of 1.1 A_260_/ml (Upa1-Cbf5) and 1.5 A_260_/ml (Rbp95-Cbf5). Samples (3.5 µl per grid) were applied to glow-discharged R3/3 holey copper grids with a 2 nm continuous carbon support film (Quantifoil), using a Vitrobot Mark IV (FEI Company) operated at 4 °C and 95% humidity. The samples were incubated on the grids for 45 seconds, the excess sample was blotted for three seconds, and the grids were plunged directly into liquid ethane. Cryo-EM data were collected on a Titan Krios G3 (FEI Company) operating at 300 kV, equipped with a Falcon 4i direct detector and a Selectris X energy filter (Thermo Fisher Scientific) under low-dose conditions (40 frames per movie, with a total dose of 40 e^-^ /Å^2^). Data were collected with a nominal pixel size of 0.727 Å/pixel and a defocus range of -0.5 to 3.5 µm, using EPU (Thermo Fisher Scientific). MotionCor2^76^ was used for motion correction of the collected movies and contrast-transfer function (CTF) parameters estimation was performed with CTFFIND4^77^. Due to the particles’ preferential orientations, both datasets were collected at 0° and 25° tilt. For the Upa1-Cbf5 dataset, 14,629 micrographs at 0° and 14,442 micrographs at 25° were collected, and for the Rbp95-Cbf5 dataset, 13,387 micrographs at 0° and 5,795 micrographs at 25° were collected.

### Cryo-EM data processing

For the Upa1-Cbf5 dataset, 14,019 micrographs at 0° tilt and 12,264 micrographs at 25° were selected for data processing, as were 12,149 micrographs at 0° and 4,041 micrographs at 25° for the Rbp95-Cbf5 sample. The 0° and 25° micrographs were combined and imported into cryoSPARC^78^ for each dataset. Patch CTF estimation was then performed, after which the initial particles were picked using the Blob Picker tool with a diameter range of 140-200 Å. A total of 5,355,748 particles were extracted for Upa1-Cbf5 and 3,227,424 for Rbp95-Cbf5, with a pixel size of 1.454 Å and a box size of 200×200 pixel. These particles were subsequently used for 2D classification and good 2D class averages (547,625 particles for Upa1-Cbf5 and 269,617 particles for Rbp95-Cbf5) were selected for *ab initio* reconstruction and non-uniform refinement. To obtain more particles, especially those with rare views, the particles were combined with those from poor 2D class averages. After three consecutive rounds of heterogeneous refinement, a snR37 snoRNP class was obtained with 814,909 particles for the Upa1-Cbf5 dataset and, respectively, with 368,472 particles for the Rbp95-Cbf5 dataset. These particles were extracted at a pixel size of 0.727 Å/pixel (box size 400×400 pixel) and were then used for non-uniform refinement and subsequent 3D classification or 3D variability analysis. As the two reconstructions were highly similar, we combined the particles obtained after heterogeneous refinement of both datasets, extracted them with a pixel size of 0.727 Å (box size 400×400 pixel), and performed non-uniform refinement and 3D classification. The final class contained 310,233 particles, achieving an overall resolution of 2.77 Å. Local filtering was performed on the map, and local refinement and local filtering were performed on the 5’ and 3′ half of the snR37 snoRNP (**Figures S7** A-F). A composite map was created in ChimeraX^79^ using the vop max command. The cryo-EM processing scheme is illustrated in detail in **Figure S6**.

### Model building

The molecular model of the snR30 snoRNP (PDB-ID: 9G25)^22^ was rigid-body fitted and used as an initial model for the H/ACA core proteins (Cbf5, Nop10, Nhp2). For the 5′ Gar1 copy and the Upa1-Upa2 heterodimer AlphaFold^80^ and AlphaFold-Multimer^81^ models were rigid-body fitted. All models were manually adjusted with Coot.^82^ For the snR37 model, the conserved regions, including the H and ACA box, of the snR30 snoRNP were used as an initial model, and the snR37 snoRNA was built *de novo* in Coot according to the cryo-EM density and assisted by secondary structure predictions. The final model was real-space refined with Phenix (**Table S3**).^83^ Cryo-EM densities and the molecular model were visualized using ChimeraX.

### Hydroxyl radical footprinting

Wild-type and *snr37*Δ mutant yeast strains were grown in YPD medium, plasmid-containing *snr37*Δ strains were grown in SDC-Trp medium. For each footprinting experiment, 10 ml cultures were grown at 30 °C to an OD_600_ of 2. Cells were rapidly chilled, harvested by centrifugation, washed with 1X PBS buffer, and resuspended in 300 µl of TM buffer (10 mM Tris-HCl pH 7.5, 1 mM MgCl_2_). Aliquots (5 µl) were transferred to 0.2 ml PCR tubes, snap-frozen in liquid nitrogen, and stored at -80 °C.

Frozen samples were irradiated for 150 ms at a synchrotron X-ray beam using a pre-chilled multi-sample holder mounted on a motorized stage (beamline 17-BM XFP, NSLS-II, Brookhaven National Laboratory, NY, USA).^84^ The ring current was 500 mA, and exposure time was controlled using a shutter actuator. When not mounted in the sample holder, samples were kept frozen on dry ice throughout the experiment. Irradiated samples were compared with control samples that were placed in the sample holder but not exposed to the beam.

Total RNA was purified using the hot acid phenol method^85^. Irradiated cells were thawed and pelleted by centrifugation. Pellets were resuspended in 400 µl TES buffer (10 mM Tris-HCl pH 7.5, 10 mM EDTA, 0.5% SDS) and mixed with 400 µl acid phenol-chloroform-isoamyl alcohol (25:24:1). Samples were vortexed for 10 s and incubated at 65 °C for 45 min with vortexing every 10-15 min. Samples were then chilled on ice for 5 min and centrifuged at maximum speed for 5 min at 4 °C. This was followed by an additional phenol-chloroform-isoamyl alcohol (25:24:1) extraction and one chloroform-isoamyl alcohol (24:1) extraction. RNA was precipitated by adding 40 µl of 3 M sodium acetate (pH 5.2) and 1 ml of 100% ethanol. RNA pellets were resuspended in nuclease-free water.

For DNase treatment, 25 µg of total RNA was incubated with 5 U DNase I (Thermo Scientific) in a total volume of 50 µl for 20 min at RT, followed by cleanup using the Zymo Research RNA Clean & Concentrator-25 kit.

For primer extension analysis, 1-3 µg of total RNA was reverse-transcribed using SuperScript III (Invitrogen) and a ^32^P-labeled primer (25S: 5′-CCAATTATCCGAATGAACTGTTCCTCTCG-3′; 18S: 5′-GCCTGCTGCCTTCCTTGGATG-3′). Sequencing ladders were generated using the same primer with a 1:1 molar ratio of ddNTPs to dNTPs. RNA was degraded with NaOH, and cDNA was purified by ethanol precipitation.^86^ Samples were resuspended in 4 µl loading dye (95% deionized formamide, 0.025% (w/v) bromophenol blue, 0.025% (w/v) xylene cyanol FF, 5 mM EDTA pH 8.0) and resolved on 6% denaturing urea-PAGE sequencing gels. Gels were run at 75 W in 1x TBE buffer and imaged using a GE Typhoon 5 system. Band intensities were quantified using SAFA,^87^ normalized to the full-length product in each lane, and plotted using GraphPad Prism.

## Data Availability

Cryo-EM reconstructions of the snR37 H/ACA snoRNP were deposited to the Electron Microscopy Data Bank (EMDB) with the accession codes EMD-XXXXX (Composite map), EMD-XXXXX (Consensus map), EMD-XXXXX (5′ half local refinement), and EMD-XXXXX (3′ half local refinement). The molecular model of the snR37 H/ACA snoRNP was deposited to the Protein Data Bank (PDB) with the accession code pdb_XXXX. The mass spectrometry data were deposited to the ProteomeXchange Consortium (www.proteomexchange.org) via the Proteomics Identification Database (PRIDE^88^) partner repository with the dataset identifier PXD075098 for the MS of excised gel bands, and with the dataset identifier PXD075805 for the TurboID data. NGS analysis data from the CRAC experiment were deposited in the Gene Expression Omnibus database under the accession number GSE324330.

## Acknowledgments

We thank Otto Berninghausen, Charlotte Ungewickel, and Susanne Rieder for cryo-EM data collection. We thank Herbert Tschochner, Sabine Rospert, and Jonathan Warner for the generous gift of antibodies and Jesús de la Cruz for the generous gift of plasmids.

## Author contributions

Conceptualization, J. H., M. T., F. D., R. B., A. K. H., B. P.; Investigation, J. H., M. T., H. Hamze, A. F., I. A. D., H. Hebbachi, M. K., A. H., S. F., S. L., S. I., T. D., K. Schlick, T. F., P. B., K. Schindlmaier, S. S., Y. H., A. K. H.; Formal analysis: J. H., M. T., M. K., S. F., S. L., T. D.; Validation, J. H., M. T., H. Hamze, M. K.; Visualization, J. H., M. T.; Supervision, E. H., D. K., Y. H., S. W., F. D., R. B., A. K. H., B. P.; Funding acquisition, D. K., S.A.W., F. D., R. B., A. K. H., Y. H., B. P.; Writing – original draft: J. H., M. T., B. P.; Writing – review & editing: J. H., M. T., H. Hamze, I. A. D., H. Hebbachi, M. K., S. F., S. I., K. Schindlmaier, E. H., D. K., Y. H., S. W., F. D., R. B., A. K. H., B. P.

