## Supplementary Material for "A bifunctional H/ACA snoRNP mediates both pseudouridylation and rRNA scaffolding during ribosome assembly"

3 BioTechMed-Graz, Graz, Austria.

4 Gene Center, University of Munich, Feodor-Lynen-Straße 25, 81377 Munich, Germany.

5 Molecular, Cellular and Developmental Biology Unit (MCD), Centre de Biologie Intégrative (CBI), Université de Toulouse, CNRS, UPS, 31062, Toulouse, France.

6 Département des sciences biologiques, Université du Québec à Montréal, 141 Av. du Président-Kennedy, Montréal, QC, H2X 1Y4, Canada

7 Centre d'excellence en recherche sur les maladies orphelines – Fondation Courtois (CERMO-FC), Université du Québec à Montréal, 141 Av. du Président-Kennedy, Montréal, QC, H2X 1Y4, Canada

8 T. C. Jenkins Department of Biophysics, Johns Hopkins University, 3400 N. Charles St., Baltimore, MD 21218, United States.

9 Department of Biology, University of Fribourg, Chemin du Musée 10, 1700 Fribourg, Switzerland.

10 Biochemistry Center, University of Heidelberg, Im Neuenheimer Feld 328, 69120 Heidelberg, Germany

11 Center for Synchrotron Biosciences, Case Western University, School of Medicine, 10900 Euclid Avenue, Cleveland, OH 44106, USA

12 These authors contributed equally

13 Lead contact

**Keywords:** H/ACA snoRNP, snoRNA, snR37, RNA folding, ribosome biogenesis, Npa1 complex, yeast

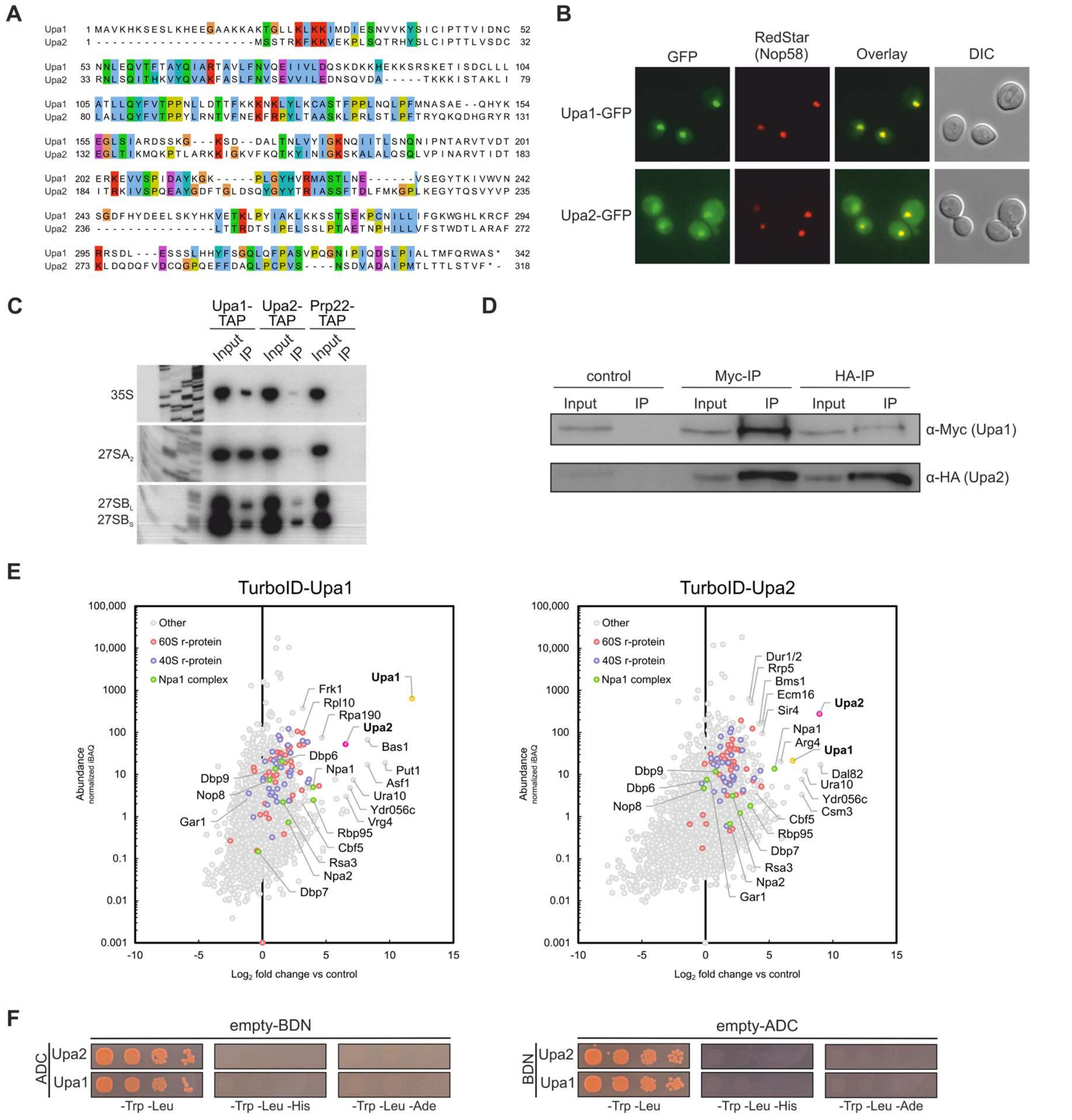


**Figure S1**. **Upa1 and Upa2 are two novel ribosome biogenesis factors.**

(**A**) Protein sequence alignment of Upa1 and Upa2. Sequences were aligned with Clustal Omega and viewed in Jalview. (**B**) Fluorescence microscopy reveals that Upa1-GFP and Upa2-GFP are localized in the nucleolus. Nop58-RedStar was used as a nucleolar marker. Overlay shows co-localization of Nop58-RedStar with Upa1-GFP and Upa2-GFP. DIC, differential interference contrast. Due to a weaker fluorescence signal, the image of Upa2-GFP was processed differently than the Upa1-GFP image to better visualize the nucleolar localization of Upa2-GFP. (**C**) Primer extension experiments suggest that Upa1-TAP and Upa2-TAP co-purify the early pre-rRNAs 35S, $27SA_2$, and 27SB. (**D**) Immunoprecipitation of Upa1-Myc or Upa2-HA followed by western blotting. (**E**) TurboID-based proximity labeling assays with the TurboID-Upa1 and TurboID-Upa2 baits. Detected proteins are plotted by normalized abundance (iBAQ, intensity-based absolute quantification; y-axis) and relative enrichment ($log_2$-transformed fold change; x-axis) compared to a control (SV40NLS-TurboID-yEGFP). Enriched proteins appear on the right of the plot. (**F**) Negative controls of Y2H interaction assays between Upa1, Upa2, and Rbp95 shown in **Figure 1**D and **1**F. Cells expressing the indicated proteins fused to AD or BD in combination with an empty vector (empty-BDN, empty-ADC) were tested as negative controls. Transformed cells were spotted in a serial dilution on SDC-Trp-Leu (growth control), SDC-Trp-Leu-His (growth on these plates indicates weak interaction), and SDC-Trp-Leu-Ade (growth on these plates indicates strong interaction).

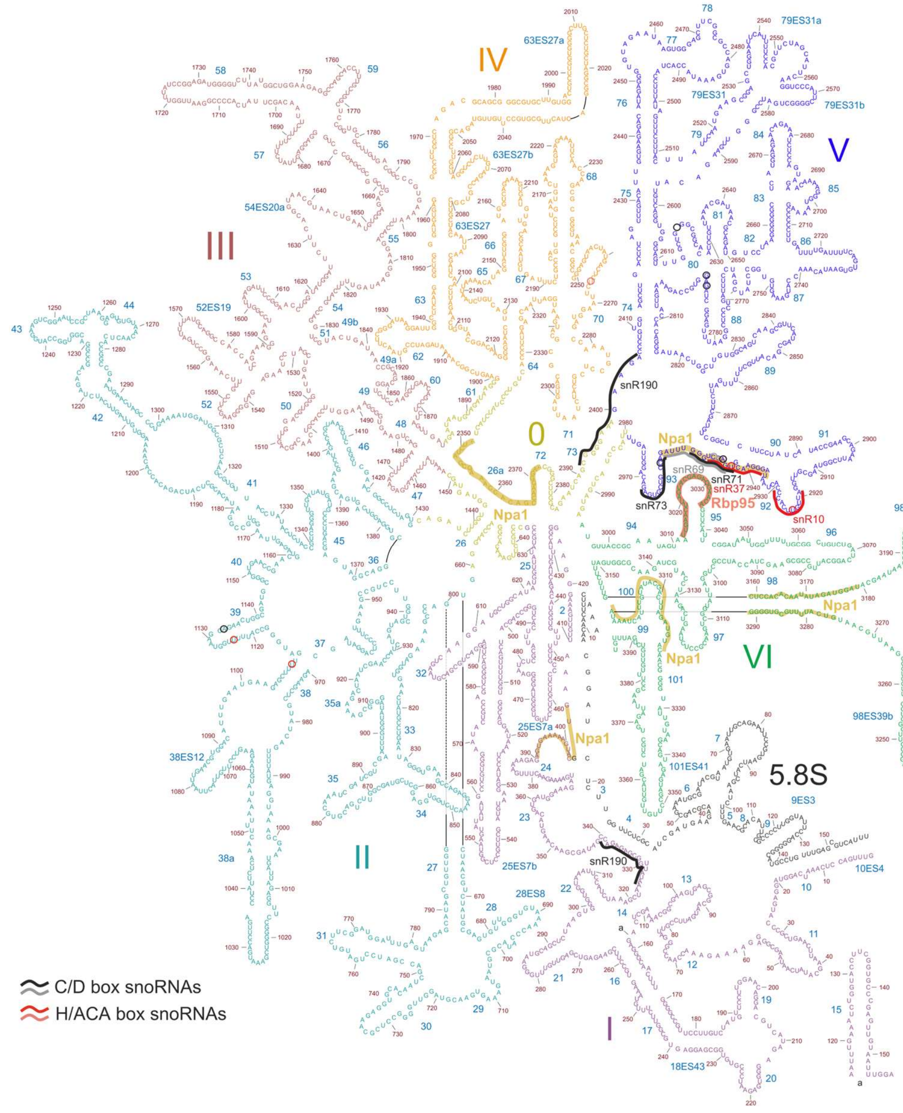


**Figure S2. 25S rRNA secondary structure and binding site of snR37.**

Secondary structure of 25S and 5.8S rRNAs. Binding sites of Rbp95[1], Npa1[2], and snoRNAs snR37, snR69, snR71, and snR73[3] are indicated.

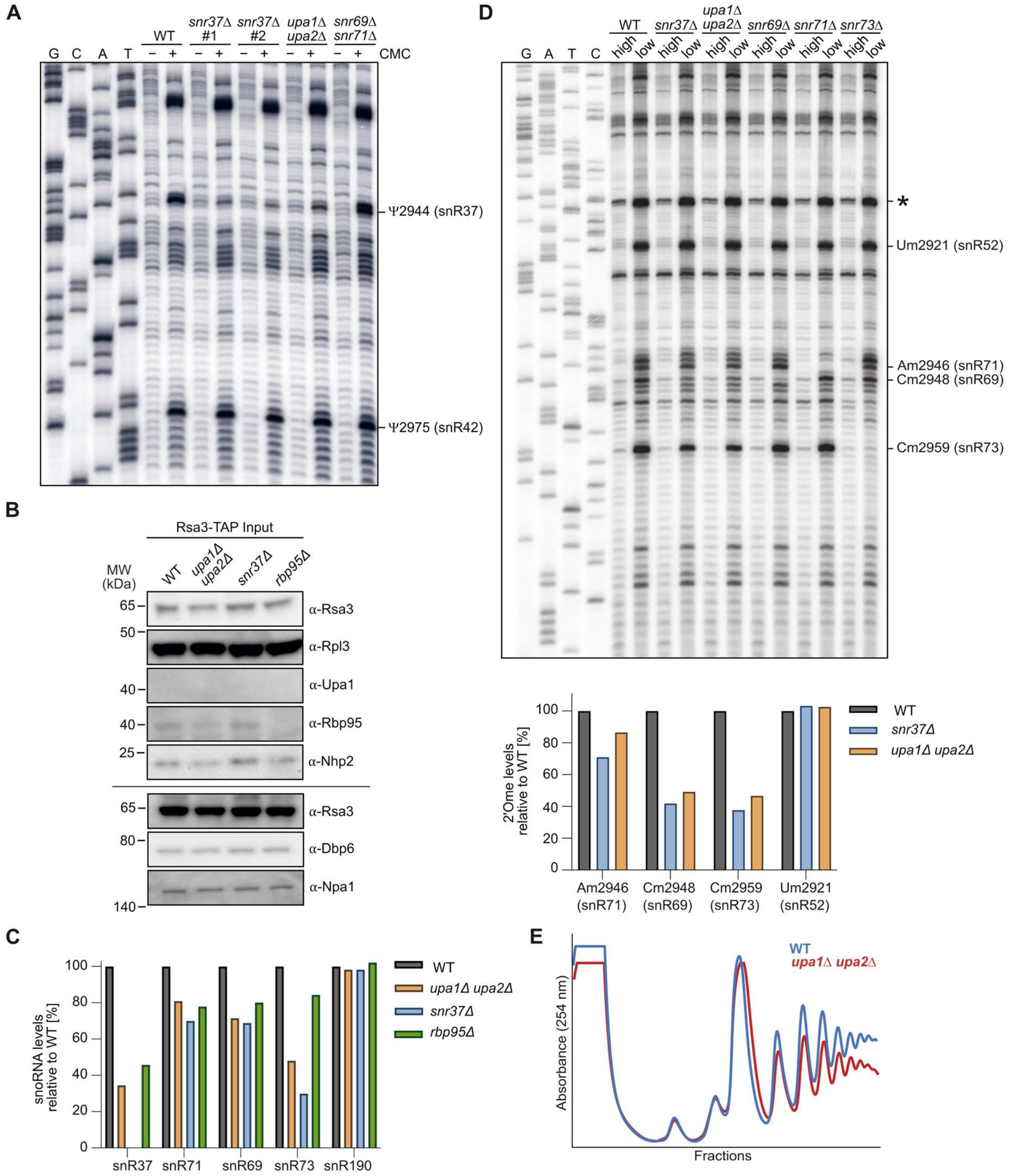


**Figure S3. Knockout of *UPA1* and *UPA2* leads to alterations in early pre-60S particles.**

(**A**) Full gel image of the CMC primer extension section displayed in **Figure 2**D. (**B**) Western blot analysis of cell lysates used as inputs for the two-step affinity purification via the Rsa3-TAP bait protein (shown in **Figure 2E**). Blots were probed with the indicated antibodies. The upper five panels are derived from a single membrane, whereas the lower three panels originate from the same samples resolved on a separate gel. Rsa3 was detected on both membranes to enable comparison between the two blots. (**C**) Quantification of northern blot data shown in **Figure 2**F (Rsa3-TAP, right panel). Levels of all snoRNAs were normalized to the snR3 signal, and the WT ratios were set to 100%. (**D**) Primer extension experiments to detect changes in rRNA methylation pattern in wild-type cells compared to *upa1*Δ *upa2*Δ or different snoRNA deletion mutants. Total RNA isolated from wild-type yeast cells (WT) and *snr37*Δ, *upa1*Δ *upa2*Δ, *snr69*Δ, *snr71*Δ, and *snr73*Δ mutant strains was analyzed by primer extension to detect methylation of rRNA. 2′-*O*-ribose methylation (2′Ome) leads to a reverse transcriptase stop under low dNTP conditions. High dNTP conditions serve as control. For the quantification shown below, a primer extension stop occurring in all samples, labeled by an asterisk (*), was used for normalization and the WT ratios were set to 100%. (**E**) Comparison of polysome profiles of wild-type (WT) and *upa1*Δ *upa2*Δ cells.

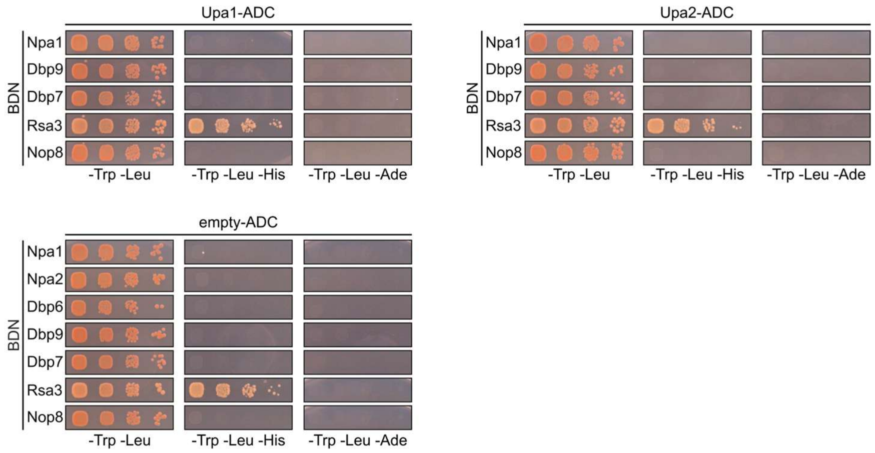


**Figure S4. Negative controls of Y2H assays between Upa1 and Upa2 and Npa1 complex members.**

For Y2H assays, cells were spotted in a serial dilution on SDC-Trp-Leu, SDC-Trp-Leu-His, and SDC-Trp-Leu-Ade plates. Proteins of interest were either fused C-terminally to the Gal4 activation domain (ADC) or fused N-terminally to the Gal4 DNA-binding domain (BDN). Note that BDN-Rsa3 resulted in growth on SDC-Trp-Leu-His plates in all combinations, including the negative control with empty-ADC, indicating self-activation and no true interaction.

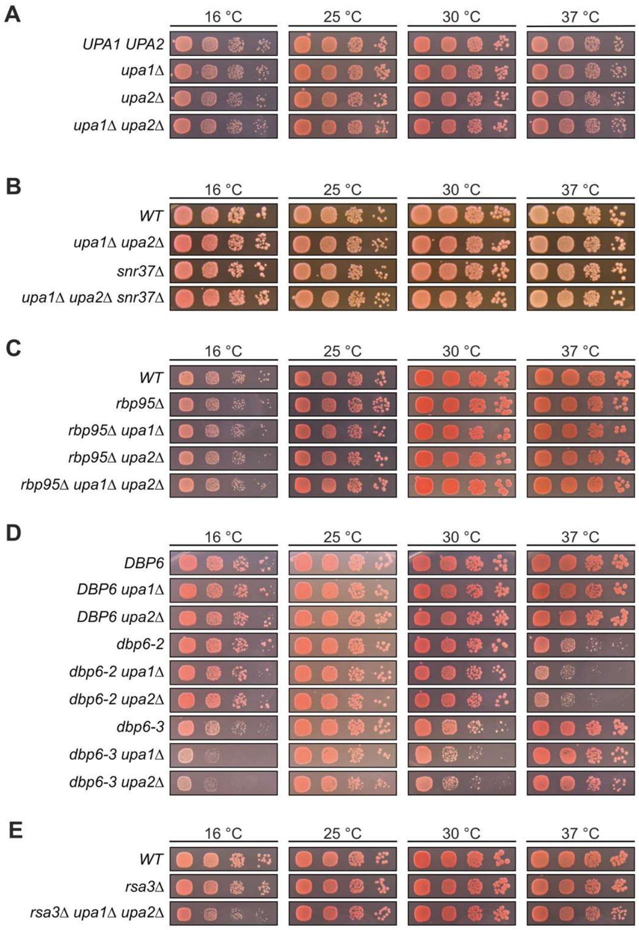


**Figure S5. Upa1 and Upa2 are functionally linked to the Npa1 complex.**

(**A-C, E**) Growth phenotypes of the indicated single and combined mutants spotted in serial dilutions on SDC plates, which were incubated for 3 days (25, 30, and 37 °C) or 6 days (16 °C). (**D**) *DBP6* deletion strains in combination with single knockouts of *UPA1* or *UPA2* carrying *LEU2* plasmids with either wild-type or different mutant alleles of *DBP6* were spotted on SDC-Leu plates, which were incubated for 3 days (25, 30, and 37 °C) or 6 days (16 °C).

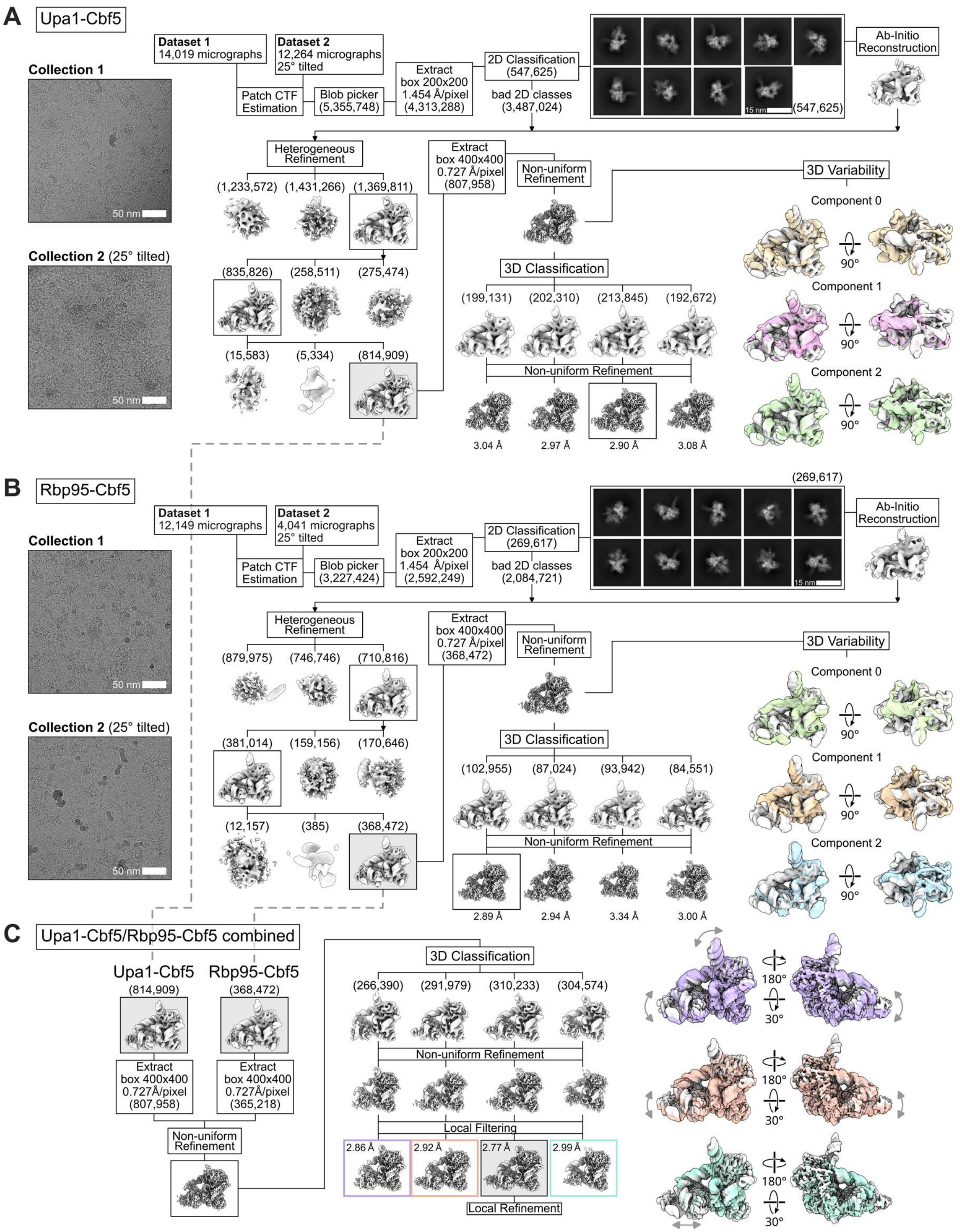


**Figure S6. Cryo-EM sorting schemes.**

(**A**, **B**) Sorting schemes for the Upa1-Cbf5 (A) and Rbp95-Cbf5 (B) datasets are shown. Representative electron micrographs (scale bar: 50 nm) are shown on the left and the processing schemes are illustrated. The initial 2D class averages are shown (scale bar: 15 nm), and the number of individual particles is indicated in parentheses. For 3D variability analysis, the first and last frame for each component was overlaid and displayed (right panels). (**C**) Classes after heterogeneous refinement for the Upa1-Cbf5 and the Rbp95-Cbf5 datasets were combined and used for 3D classification. The final class resolved with an overall resolution of 2.77 Å was used for local refinement and model building. The cryo-EM density maps of the other classes were overlaid with the final class and are shown on the right.

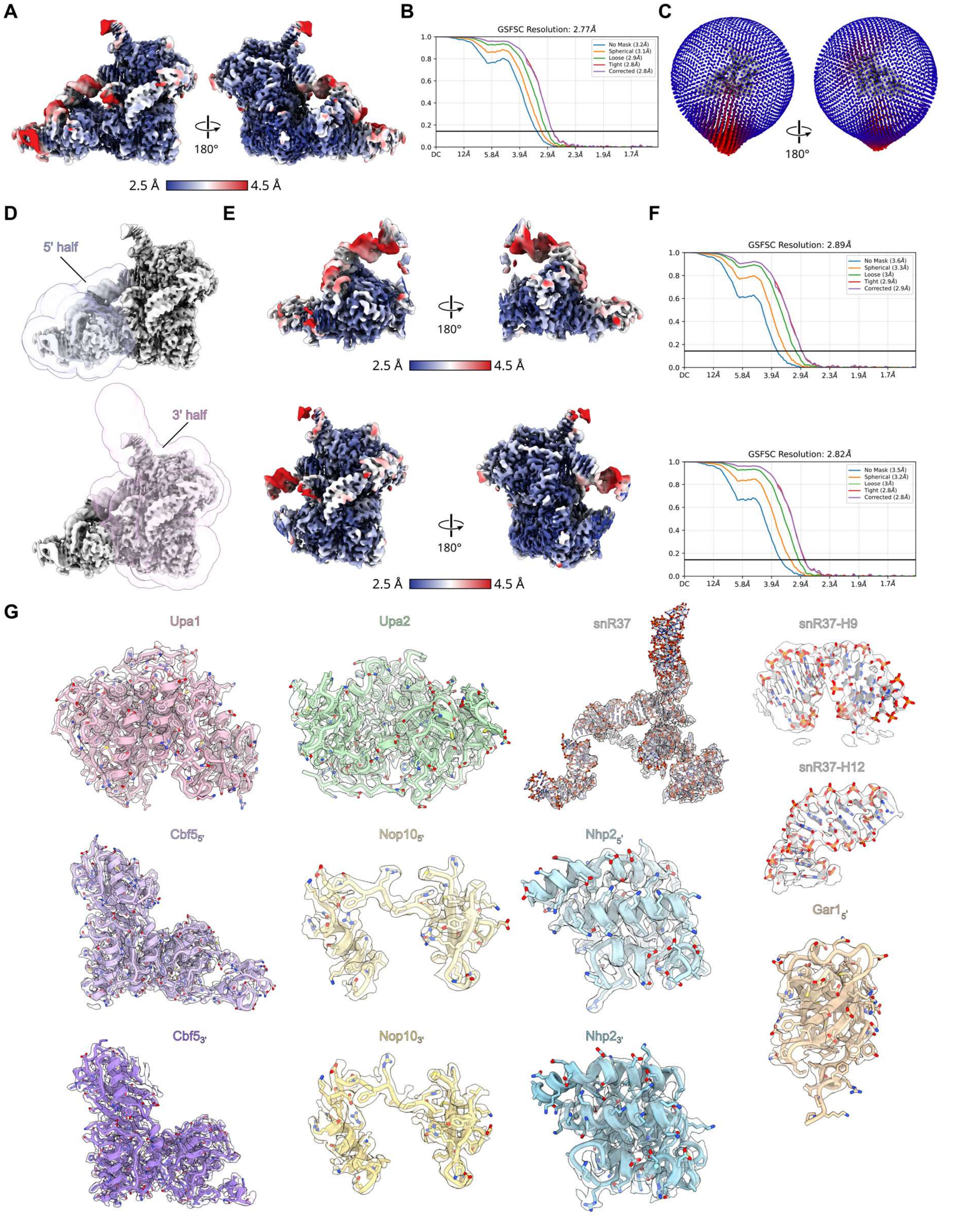


**Figure S7. Local resolution, FSC curves, and cryo-EM densities and models of the snR37 snoRNP.**

(**A**-**C**) Local resolution filtered density map (A), Fourier shell correlation (FSC) curve (B), and 3D representation of the angular distribution (C) of the snR37 snoRNP. (**D**-**F**) Local refinements of the 5′ and 3′ modules of the snR37 snoRNP. The applied masks (D), density maps colored according to local resolution (E), and FSC curves (F) are shown. (**G**) Molecular models of the individual snR37 snoRNP components with overlaid segmented cryo-EM density maps.

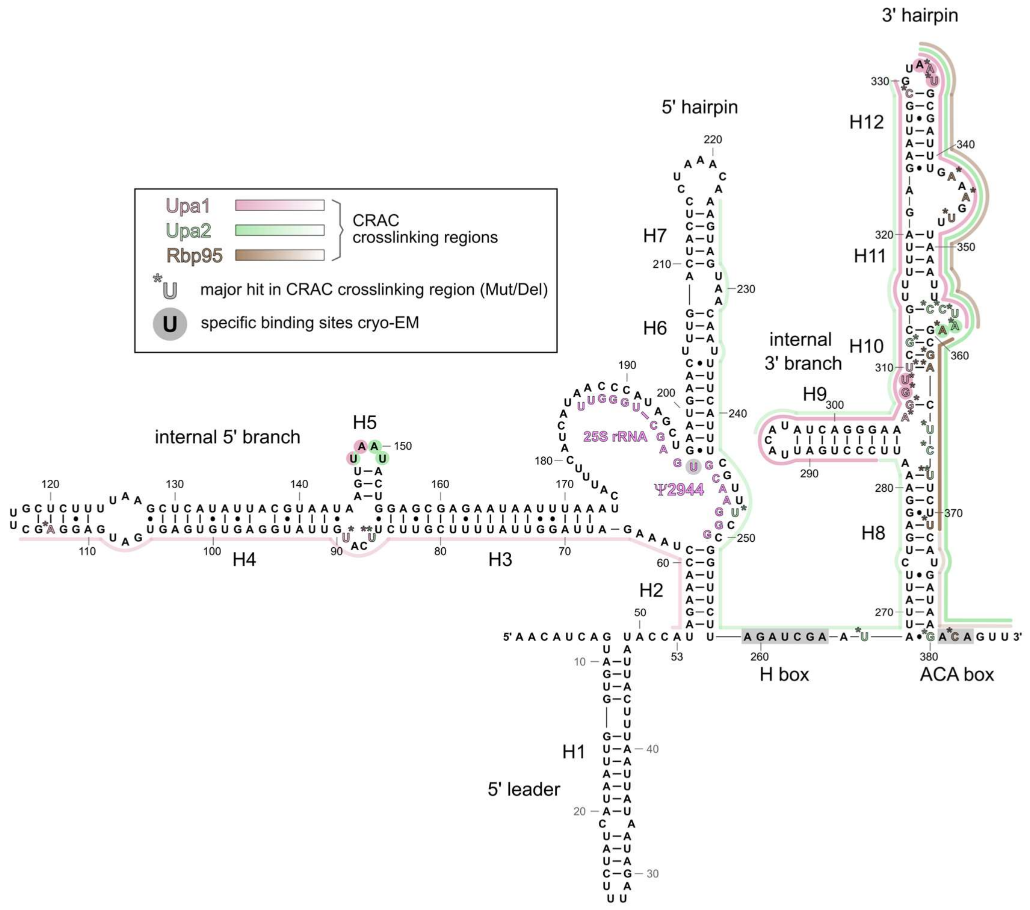


**Figure S8: Secondary structure of the snR37 snoRNA.**

Structure-based secondary structure of the snR37 snoRNA. Binding sites of Upa1 and Upa2, as observed in the cryo-EM structure, are indicated with pink and light green circles, respectively. Major sequencing hits (colored line next to snR37sequence) and major deletion/mutation hits (colored letters with an asterisk (*)) found in the CRAC experiment are indicated. The part of 25S rRNA domain V that base-pairs with the pseudouridylation pocket of the 5′ hairpin is indicated by magenta letters and the modification site (Y2944) is highlighted. The different regions of snR37 are labeled.

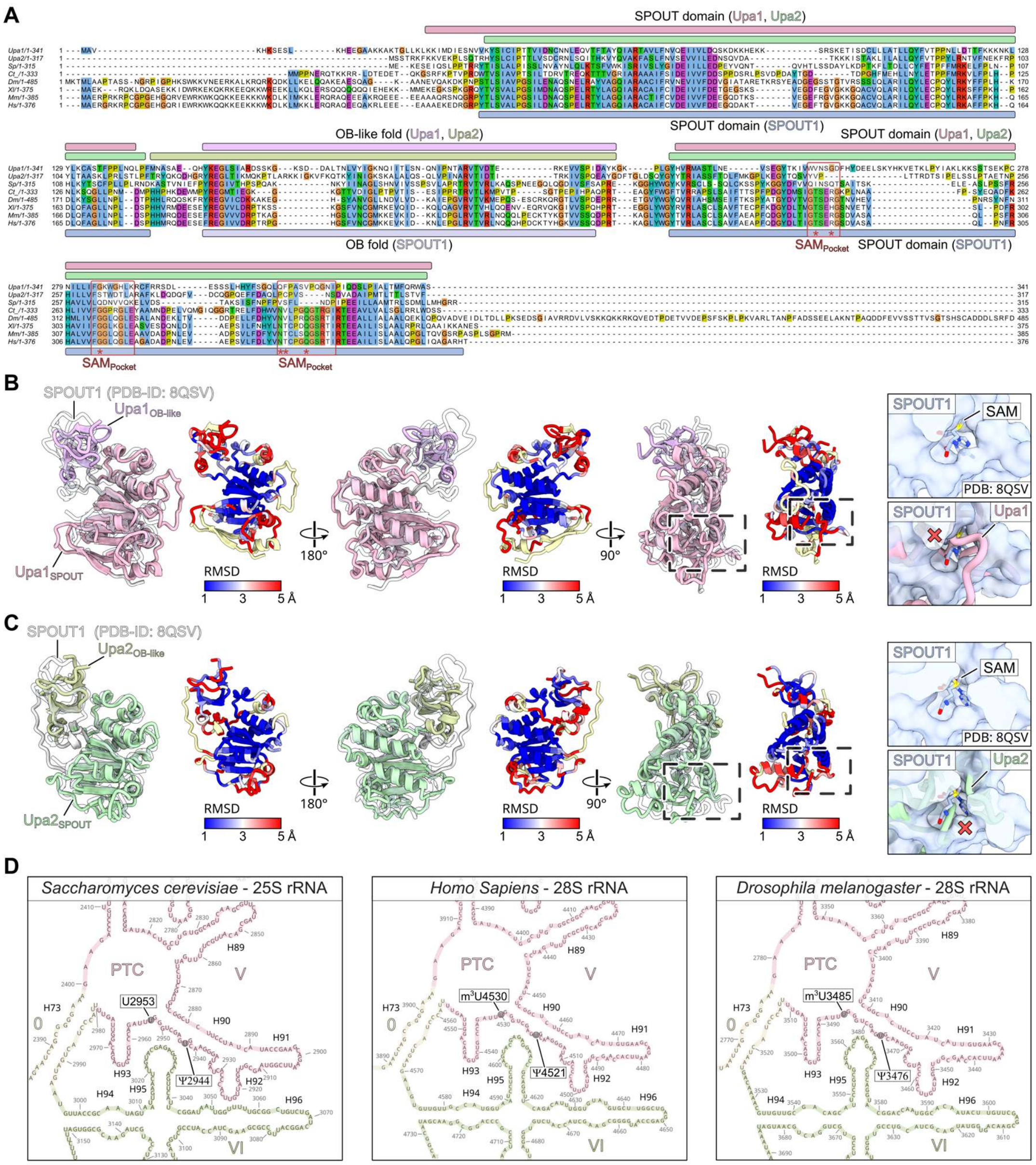


**Figure S9. Comparison of Upa1-Upa2 and human SPOUT1.**

(**A**) Sequence alignment of Upa1 and Upa2 and their homologs from *Schizosaccharomyces pombe* (*Sp*), *Chaetomium thermophilum* (*Ct*), *Drosophila melanogaster* (*Dm*), *Xenopus laevis* (*Xl*), *Mus musculus* (*Mm*), and *Homo sapiens* (*Hs*). The sequences were aligned using Clustal Omega and displayed with Jalview. The SPOUT and OB-like fold domain boundaries for Upa1 and Upa2 are indicated above the alignment. For the human SPOUT1 homolog, the SPOUT domain and OB fold are shown below the alignment, and the residues that contribute to the SAM-binding pocket (SAM$_{Pocket}$) within SPOUT1 are highlighted with red boxes. Residues that directly interact with SAM are marked with asterisks (*). (**B**, **C**) Upa1 and Upa2 in different orientations. Molecular models of Upa1 (B) and Upa2 (C) with labeled SPOUT and OB-like domains and overlaid with a model of human SPOUT1 (PDB-ID: 8QSV) [4] (left panels). The same orientations are shown, colored by RMSD based on structural alignment with SPOUT1 (right panels). The region of the SAM-binding pocket within SPOUT1 is indicated with dashed boxes. A close-up of the surface view of SPOUT1 bound to SAM is shown in the upper right panels, along with the superposition of the molecular models of either Upa1 (B) or Upa2 (C). (**D**) Close-ups of the secondary structures of the large subunit rRNA from *S. cerevisiae*, *H. sapiens*, and *D. melanogaster* (modified from http://apollo.chemistry.gatech.edu/RibosomeGallery/). Residue Ψ2944, which is modified by the snR37 snoRNP, the equivalent residue in *H. sapiens* (Ψ4521) and *D. melanogaster* (Ψ3476), and the residue modified by human SPOUT1 (m$^3$U4530) and fly Ptch (m$^3$U3485) are labeled, as is the equivalent nucleotide in *S. cerevisiae* (U2953). For *D. melanogaster* the nucleotide numbering according to [5] was used.

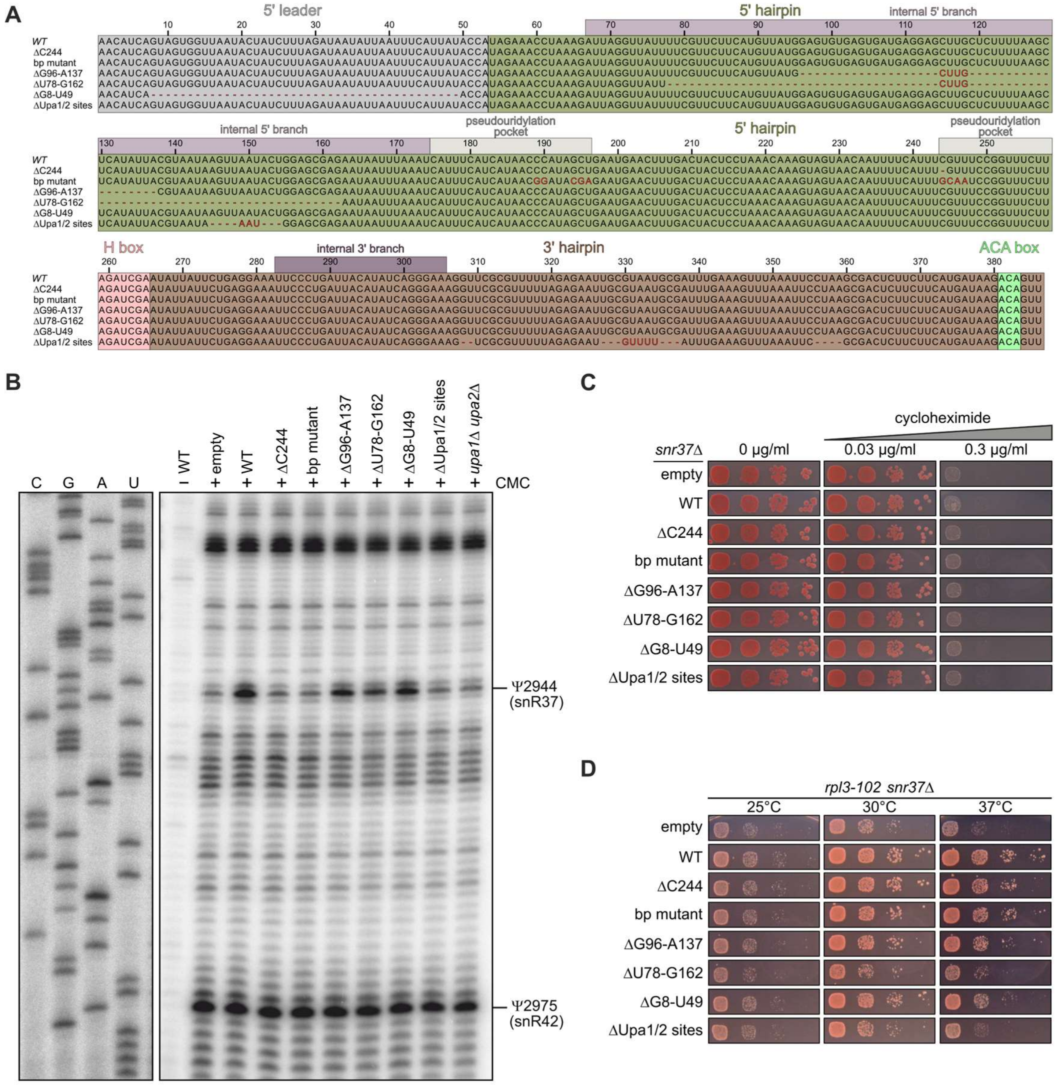


**Figure S10. Upa1 and Upa2 are required for pre-60S recruitment of the snR37 snoRNP.**

(**A**) Sequence alignment of the utilized snR37 wild-type (WT) and mutant constructs. Mutations are shown in red and 5′ leader, 5′ and 3′ hairpin, and H and ACA box consensus sequences are highlighted. (**B**) Full gel image of the CMC primer extension section displayed in **Figure 7**C. (**C**) Control experiment for the experiment shown in **Figure 7**D, indicating that none of the *snr37* mutants exhibits resistance to the translation inhibitor cycloheximide. An *snr37*Δ strain was transformed with *TRP1* plasmids expressing the indicated snR37 variants and cells were spotted in serial dilutions on SDC-Trp plates containing different concentrations of cycloheximide (0, 0.03, or 0.3 µg/ml). Plates were incubated at 30 °C for 2 days. (**D**) Growth complementation assay. An *rpl3-102 snr37*Δ double mutant strain was transformed with *TRP1* plasmids expressing the indicated snR37 variants and cells were spotted in serial dilutions on SDC-Trp plates, which were incubated for 2 days at the indicated temperatures.

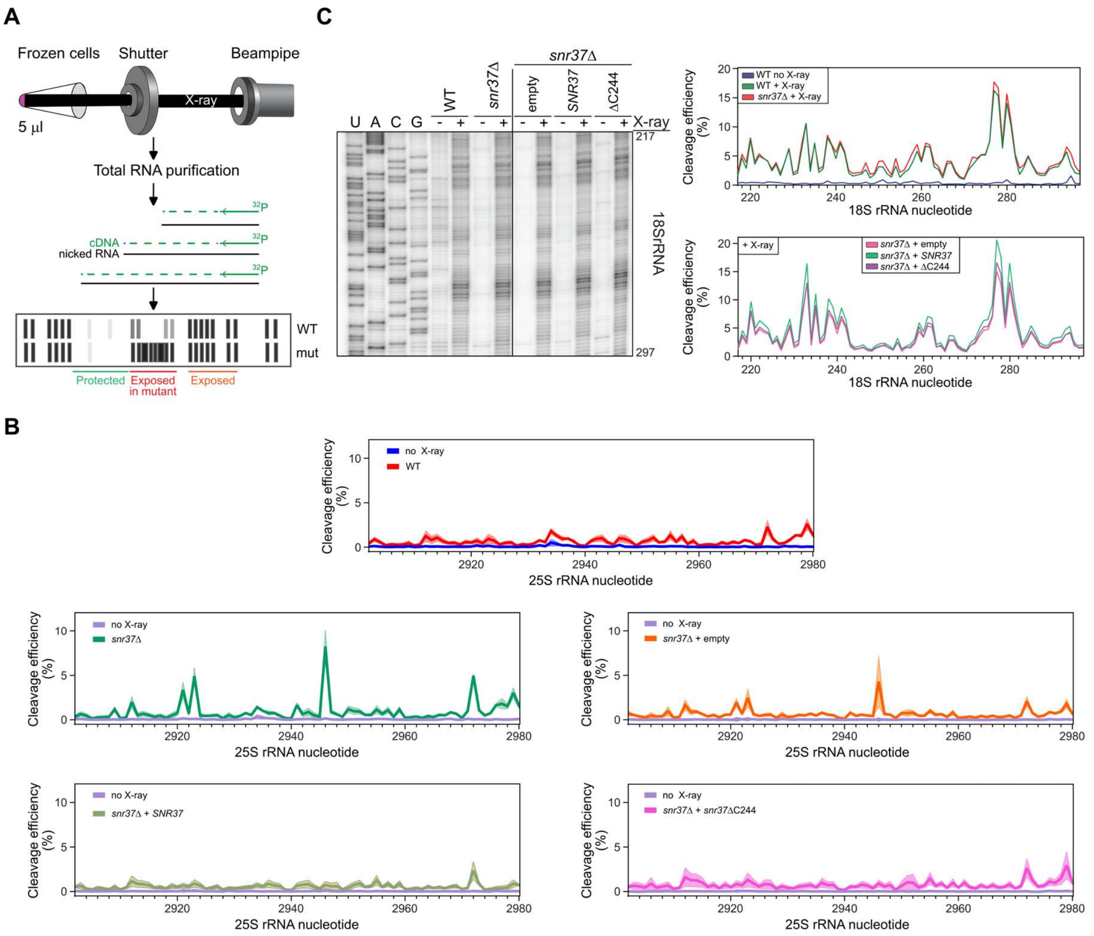


**Figure S11. 25S but not 18S rRNA is perturbed in *snr37*Δ strains.**

**(A)** Alignment of frozen aliquoted samples to the X-ray beamline and downstream sample analysis. **(B)** Normalized cleavage efficiencies of individual nucleotides in 25S rRNA, determined by *in vivo* X-ray hydroxyl radical footprinting, are shown for WT (red) and *snr37*Δ (teal) strains, as well as for *snr37*Δ strains transformed with *TRP1* plasmids harboring no insert (empty; orange), *SNR37* (pale green), or *snr37*ΔC244 (pink). Light shades indicate the range from two biological replicates. Plots include corresponding no-irradiation controls. **(C)** Sequencing gel and cleavage efficiency plots of 18S rRNA after 150 ms X-ray exposure in the same strains. Cleavage profiles were unchanged among strains.

### Supplementary Tables

**Table S1. Yeast strains**

| name | genotype | source |
|---|---|---|
| W303 | *ade2-1 his3-11,15 leu2-3,112 trp1-1 ura3-1 can1-100* | [6] |
| C303 | *his3-11,15 leu2-3,112 trp1-1 ura3-1 can1-100* | [7] |
| Upa1-GFP Nop58-RedStar | C303 *MAT*α *UPA1*-GFP::kanMX4 *NOP58*-RedStar::natNT2 | this study |
| Upa2-GFP Nop58-RedStar | C303 *MAT*α *UPA2*-GFP::kanMX4 *NOP58*-RedStar::natNT2 | this study |
| Y2H PJ69-4A | *MAT***a** *trp1-901 leu2-3,112 ura3-52 his3-200 gal4*Δ *gal80*Δ *LYS2*::*GAL1-His3 GAL2-Ade2 met2*::*GAL7-lacZ* | [8] |
| *upa1*Δ | W303 *MAT***a** *upa1*Δ:: klTRP1 | this study |
| *upa2*Δ | W303 *MAT***a** *upa2*Δ:: kanMX | this study |
| *upa1*Δ *upa2*Δ | W303 *MAT***a** *upa1*Δ:: klTRP1 *upa2*Δ::kanMX | this study |
| *snr37*Δ | W303 *MAT***a** *snr37*Δ:: natNT2 | this study |
| *snr69*Δ *snr71*Δ | W303 *MAT***a** *snr69*Δ::HIS3MX *snr71*Δ::hphNt1 | this study |
| *snr69*Δ | W303 *MAT***a** *snr69*Δ::*HIS3MX* | this study |
| *snr71*Δ | W303 *MAT***a** *snr71*Δ::hphNt1 | this study |
| *snr73*Δ | W303 *MAT***a** *snr73*Δ:: HIS3MX | this study |
| *upa1*Δ *upa2*Δ *snr37*Δ | W303 *MAT***a** *upa1*Δ::klTRP1 *upa2*Δ::kanMX *snr37*Δ::natNT2 | this study |
| *rbp95*Δ | W303 *MAT***a** *rbp95*Δ::natNT2 | [1] |
| *rbp95*Δ *upa1*Δ | W303 *MAT***a** *rbp95*Δ::natNT2 *upa1*Δ::klTRP1 | this study |
| *rbp95*Δ *upa2*Δ | W303 *MAT***a** *rbp95*Δ::natNT2 *upa2*Δ::kanMX | this study |
| *rbp95*Δ *upa1*Δ *upa2*Δ | W303 *MAT***a** *upa1*Δ::klTRP1 *upa2*Δ::kanMX *rbp95*Δ::natNT2 | this study |
| *rbp95*Δ *snr37*Δ | W303 *MAT***a** *rbp95*Δ::kanMX *snr37*Δ::natNT2 | this study |
| *rsa3*Δ | W303 *MAT***a** *rsa3*Δ::HIS3MX | this study |
| *rsa3*Δ *upa1*Δ *upa2*Δ | W303 *MAT***a** *rsa3*Δ::HIS3MX *upa1*Δ::klTRP1 *upa2*Δ::kanMX | this study |
| *DBP6* shuffle | W303 *MAT***a** *dbp6*Δ::kanMX [pRS416-*DBP6*] | [9] |
| *DBP6* shuffle *upa1*Δ | W303 *MAT***a** *dbp6*Δ::kanMX [pRS416-*DBP6*] *upa1*Δ::klTRP1 | this study |
| *DBP6* shuffle *upa2*Δ | W303 *MAT***a** *dbp6*Δ::kanMX [pRS416-*DBP6*] *upa2*Δ:: kanMX | this study |
| *DBP9* shuffle | W303 *MAT***a** *dbp9*Δ::HIS3MX [YCplac33-*DBP9*] | [10] |
| *DBP9* shuffle *upa1*Δ *upa2*Δ | W303 *MAT***a** *dbp9*Δ::HIS3MX [YCplac33-*DBP9*] *upa1*Δ::klTRP1 *upa2*Δ::kanMX | this study |
| *DBP9* shuffle *snr37*Δ | W303 *MAT***a** *dbp9*Δ::HIS3MX [YCplac33-*DBP9*] *snR37*Δ::natNT2 | this study |
| *RPL3* shuffle | W303 *MAT***a** *rpl3*Δ::HIS3MX6 [YCPlac33-*RPL3*] | [11] |
| *RPL3* shuffle *upa1*Δ *upa2*Δ | W303 *MAT***a** *rpl3*Δ::HIS3MX6 [YCPlac33-*RPL3*] *upa1*Δ::klTRP1 *upa2*Δ::kanMX | this study |

| | | |
|---|---|---|
| *RPL3* shuffle *snr37*Δ | W303 *MAT***a** *rpl3*Δ::HIS3MX6 [YCPlac33-*RPL3*] *snr37*Δ::natNT2 | this study |
| BY4741 | *MAT***a** *his3*Δ1 *leu2*Δ0 *met15*Δ0 *ura3*Δ0 | [12] |
| Upa1-HTP (for CRAC) | BY4741 *MAT***a** *UPA1*-HTP::klURA3 | this study |
| Upa2-HTP (for CRAC) | BY4741 *MAT***a** *UPA2*-HTP:: klURA3 | this study |
| Rsa3-TAP | W303 *MAT***a** *RSA3*-TAP::HIS3MX6 | this study |
| Rsa3-TAP *upa1*Δ *upa2*Δ | W303 *MAT***a** *RSA3*-TAP::HIS3MX6 *upa1*Δ::*KLTRP1* *upa2*Δ::kanMX | this study |
| Rsa3-TAP *snr37*Δ | W303 *MAT***a** *RSA3-TAP*::HIS3MX6 *snr37*Δ::natNT2 | this study |
| Rsa3-TAP *rbp95*Δ | W303 *MAT***a** *RSA3*-TAP::HIS3MX6 *rbp95*Δ::natNT2 | this study |
| Upa1-TAP | S288C *MAT***a** *his3Δ1 leu2Δ0 met15Δ0 ura3Δ0, UPA1*-TAP::HIS3MX | Open Biosystems |
| Upa2-TAP | S288C *MAT***a** *his3Δ1 leu2Δ0 met15Δ0 ura3Δ0, UPA2*-TAP::HIS3MX | Open Biosystems |
| Prp22-TAP | S288C *MAT***a** *his3Δ1 leu2Δ0 met15Δ0 ura3Δ0, PRP22*-TAP::HIS3MX | Open Biosystems |
| YPH499 | *MAT***a** *ura3-52 lys2-801_amber ade2-101_ochre trp1-Δ63 his3-Δ200 leu2-Δ1* | [13] |
| YHH1 | YPH499 *upa1*Δ::*TRP1* *upa2*Δ::*HIS3* | this study |
| YHH2 | YPH499 *snr37*Δ::kanMX6 | this study |
| YHH3 | YPH499 *upa1*Δ::*TRP1* *upa2*Δ::*HIS3* *snr37*Δ::kanMX6 | this study |
| YHH4 | YPH499 *UPA1-13MYC*::*TRP1* *UPA2-3HA*::*HIS3MX6* | this study |
| Upa1-FTpA | W303 MATα *UPA1*-FtpA::natNT2 | this study |
| Upa1-TAP Cbf5-Flag | W303 MATα *UPA1*-TAP::HIS3MX *CBF5*-Flag::natNT2 | this study |
| Rbp95-TAP Cbf5-Flag | W303 MATα *RBP95*-TAP::klURA3 *CBF5*-Flag::natNT2 | this study |
| TurboID YDK11-5A | W303 *MAT*α *ade2-1 his3-11,15 leu2-3,112 trp1-1 ura3-1 can1-100 ade3*::*kanMX4* | [9] |

**Table S2. Plasmids used in this study**

| **name** | **relevant information** | **source** |
|---|---|---|
| pFA6a-HIS3MX4 | for chromosomal deletion | [14] |
| pFA6a-kanMX4 | for chromosomal deletion | [14] |
| pFA6a-TRP1 | for chromosomal deletion | [15] |
| pFA6a-klTRP1 | for chromosomal deletion | [16] |
| pFA6a-natNT2 | for chromosomal deletion | [17] |
| pFA6a-hphNT1 | for chromosomal deletion | [17] |
| pFA6a GFP::HIS3MX4 | for C-terminal tagging | [14] |
| pFA6a TAP::HIS3MX6 | for C-terminal tagging | [18] |
| pBS1539 HTP::klURA3 | for C-terminal tagging | [19] |
| pFA6a-3HA-*HIS3*MX6 | for C-terminal tagging | [15] |
| pFA6a-13Myc-*TRP1* | for C-terminal tagging | [15] |
| pGAG4ADC111-*UPA1* | CEN, *LEU2,* P*ADH1,* T*ADH1*, C-terminal $(GA)_5$-G4AD-HA | this study |

| | | |
|---|---|---|
| pGAG4ADC111-*UPA2* | CEN, *LEU2,* P*ADH1,* T*ADH1*, C-terminal $(GA)_5$-G4AD-HA | this study |
| pG4BDC22-*RBP95* | CEN, *TRP1,* P*ADH1,* T*ADH1*, C-terminal $(GA)_5$-G4BD-cMyc | this study |
| pG4BDN22-*UPA1* | CEN, *TRP1,* P*ADH1,* T*ADH1*, N-terminal G4BD-cMyc | this study |
| pG4BDN22-*UPA2* | CEN, *TRP1,* P*ADH1,* T*ADH1*, N-terminal G4BD-cMyc | this study |
| pG4BDN22-*NPA1* | CEN, *TRP1,* P*ADH1,* T*ADH1*, N-terminal G4BD-cMyc | this study |
| pG4BDN22-*NPA2* | CEN, *TRP1,* P*ADH1,* T*ADH1*, N-terminal G4BD-cMyc | this study |
| pG4BDN22-*DBP6* | CEN, *TRP1,* P*ADH1,* T*ADH1*, N-terminal G4BD-cMyc | this study |
| pG4BDN22-*DBP9* | CEN, *TRP1,* P*ADH1,* T*ADH1*, N-terminal G4BD-cMyc | this study |
| pG4BDN22-*DBP7* | CEN, *TRP1,* P*ADH1,* T*ADH1*, N-terminal G4BD-cMyc | this study |
| pG4BDN22-*RSA3* | CEN, *TRP1,* P*ADH1,* T*ADH1*, N-terminal G4BD-cMyc | this study |
| pG4BDN22-*NOP8* | CEN, *TRP1,* P*ADH1,* T*ADH1*, N-terminal G4BD-cMyc | this study |
| pG4BDC22-*UPA1* | CEN, *TRP1,* P*ADH1,* T*ADH1*, C-terminal $(GA)_5$-G4BD-cMyc | this study |
| pG4BDC22-*UPA2* | CEN, *TRP1,* P*ADH1,* T*ADH1*, C-terminal $(GA)_5$-G4BD-cMyc | this study |
| pETDuet-1-$(His)_6$-*RBP95* | $Amp^r$, T7 promoter/*lac* operator; *RBP95* in MCS1 | [1] |
| pETDuet-1-$(His)_6$-*RBP95*/ Flag-*UPA1* | $Amp^r$, T7 promoter/*lac* operator; *RBP95* in MCS1, *UPA1* in MCS2 | this study |
| pETDuet-1-$(His)_6$-*RBP95*/ Flag-*UPA2* | $Amp^r$, T7 promoter/*lac* operator; *RBP95* in MCS1, *UPA2* in MCS2 | this study |
| pETDuet-1-Flag-*UPA1* | $Amp^r$, T7 promoter/*lac* operator; *UPA1* in MCS2 | this study |
| pETDuet-1-Flag-*UPA2* | $Amp^r$, T7 promoter/*lac* operator, *UPA2* in MCS2 | this stud |
| pETDuet-1-$(His)_6$-*UPA1*/Flag-*UPA2* | $Amp^r$, T7 promoter/*lac* operator; *UPA1* in MCS1, *UPA2* in MCS2 | this study |
| pETDuet-1-$(His)_6$-*UPA1* | $Amp^r$, T7 promoter/*lac* operator; *UPA1* in MCS1 | this study |
| pCUP111-SV40NLS-yEGFP-(GA)5-TurboID-2xHA (pDK9296) | *CEN, LEU2,* P*CUP1,* T*ADH1,* SV40NLS-*yEGFP*-TurboID-2xHA | [1] |
| pCUP111-*UPA1*-(GA)5-TurboID-2xHA (pDK9289) | CEN, *LEU2,* P*CUP1,* T*ADH1, UPA1-*TurboID-2xHA | this study |
| pCUP111-*UPA2*-(GA)5-TurboID-2xHA (pDK9288) | CEN, *LEU2,* P*CUP1,* T*ADH1, UPA2-*TurboID-2xHA | this study |
| pCUP111-SV40NLS-TurboID-(GA)5-yEGFP (pDK9934) | CEN, *LEU2,* P*CUP1,* T*ADH1,* SV40NLS-TurboID-*yEGFP* | [20] |
| pCUP111-SV40NLS-TurboID-(GA)5-*UPA1* (pDK12046) | CEN, *LEU2,* P*CUP1,* T*ADH1,* SV40NLS-TurboID-*UPA1* | this study |

| | | |
|---|---|---|
| pCUP111-SV40NLS-TurboID-(GA)5-*UPA2* (pDK12045) | CEN, *LEU2,* P*CUP1,* T*ADH1,* SV40NLS-TurboID-*UPA2* | this study |
| pRS415-*DBP6* | *CEN*, *LEU2*, P*DBP6*, *DBP6,* T*DBP6* | [21] |
| pRS415-*dbp6-2* | *CEN*, *LEU2*, P*DBP6*, *dbp6-2 (Y552C),* T*DBP6* | [9] |
| pRS415-*dbp6-3* | *CEN*, *LEU2*, P*DBP6*, *dbp6-3 (E39G S457P R463G),* T*DBP6*, | [9] |
| YCplac111-HA-*DBP9* | *CEN*, *LEU2*, P*DBP9*, HA-*DBP9*, *T*DBP9 | [22] |
| YCplac111-HA-*dbp9-5* | *CEN*, *LEU2*, P*DBP9,* HA-*dbp9-5 (F389S)*, T*DBP9* | [22] |
| YCplac111-*RPL3* | *CEN*, *LEU2*, P*RPL3*, *RPL3*, T*RPL3* | [11] |
| YCplac111-*rpl3-102* | *CEN*, *LEU2*, P*RPL3, rpl3-102 (K30E)*, T*RPL3* | [11] |

P and T denote promoter and terminator, respectively.

**Table S3. Cryo-EM data collection, refinement, and model statistics.**

| | **snR37 H/ACA snoRNP**<br>(EMD-XXXXX)<br>(pdb_XXXX) |
|---|---|
| **Data collection & processing** | |
| Camera | Falcon 4i |
| Magnification | 165,000 |
| Voltage (kV) | 300 |
| Electron exposure ($e^-/Å^2$) | 40 |
| Defocus range (μm) | 0.5 - 3.5 |
| Pixel size (Å) | 0.727 |
| Symmetry imposed | C1 |
| Micrographs collected (no.) | 42,473 |
| Initial particle images (no.) | 8,583,172 |
| Final particle images (no.) | 310,233 |
| Map resolution (Å) | 2.77 |
| FSC threshold | 0.143 |
| **Refinement** | |
| Initial model used (PDB code) | 9G25, AlphaFold2 |
| Model resolution (Å) | 2.8 |
| FSC threshold | 0.5 |
| Map sharpening B factor ($Å^2$) | -80 |
| Model composition | |
| Non-hydrogen atoms | 20,535 |
| Protein residues | 1,798 |
| Nucleotide residues | 303 |
| Ligands | 0 |
| R.m.s deviations | |
| Bond lengths (Å) | 0.005 |
| Bond angles (°) | 0.929 |
| Validation | |
| MolProbity score | 1.26 |
| Clash score | 5.00 |
| Poor rotamers (%) | 0.06 |
| Ramachandran plot | |
| Favored (%) | 98.47 |
| Allowed (%) | 1.53 |
| Disallowed (%) | 0.00 |
| Map vs. Model CC (mask) | 0.83 |
| **Local & Consensus** | EMD-XXXXX |
| **Refinements** | EMD-XXXXX |
| | EMD-XXXXX |
